# Halophytic bacterial endophytome: a potential source of beneficial microbes for a sustainable agriculture

**DOI:** 10.1101/2020.07.29.226860

**Authors:** Christos A. Christakis, Georgia Daskalogiannis, Anastasia Chatzakis, Emmanouil A. Markakis, Angeliki Sagia, Giulio Flavio Rizzo, Vittoria Catara, Ilias Lagkouvardos, David J. Studholme, Panagiotis F. Sarris

## Abstract

Halophytes have evolved several strategies to survive in saline environments; however, additional support from their associated microbiota could help combat adverse conditions. Endophytic communities of halophytes may be different than those in other plants because salinity acts as an environmental filter. At the same time, they may contribute to the host’s adaptation to adverse environmental conditions and can improve host tolerance against various biotic and abiotic stresses, which may be of importance in modern and sustainable agriculture.

In this study the culturable endophytic bacteria of three halophytic species *Cakile maritima*, *Matthiola tricuspidata* and *Crithmum maritimum* were isolated and identified. Endophytic bacteria were isolated from roots and leaves of the sampled plants. Significant differences were observed in bacterial species abundance among different plant species and tissue from which the isolates were obtained. In total, 115 strains were identified by analysis of complete 16S rDNA sequences, while the majority of these isolates were derived from the root samples.

The strains were evaluated for their ability to: 1) grow *in-vitro* in high levels of NaCl; 2) inhibit the growth of the economically important plant pathogenic fungus *Verticillium dahliae in vitro* and *in planta*, the human pathogenic fungus *Aspergillus fumigatus in vitro*, as well as, the economically important plant bacterial pathogens *Ralstonia solanacearum* and *Clavibacter michiganensis in vitro*; 3) provide salt tolerance *in planta*; 4) provide growth promoting effect *in planta*.

Additionally, the genomes of twelve selected isolates, exhibiting interesting features, were sequenced and analysed. Three novel bacterial species were identified that belong to the genus *Pseudomonas* (two strains) and *Arthrobacter* (one strain).

The outcome of our study is the proof-of-concept that the crop wild relatives (CWR) halophytic microbiome could potentially serve as a source of beneficial microorganisms that could be used (as unique species or as artificial communities) as Bio-Inoculants, for the enhancement of plant growth and stress tolerance in crops, including the high-salinity stress.

This is very important in the era of ecosystem degradation and climate change, where the maximizing microbial functions in agroecosystems could be a prerequisite for the future of global sustainable agriculture. Globally, there is a strong need for the identification and bio-banking of novel beneficial endophytic microbes with as many desirable characters, for the development of a new environmentally friendly global strategy in food production that will be based in the sustainable agriculture with low chemical inputs and a low environmental impact.

## Introduction

Bacterial endophytes are endosymbiotic microorganisms widespread among plants that colonize intercellular and intracellular spaces of all plant compartments. Each individual plant is a host to bacterial and fungal endophytes that colonize plant tissues for all or part of the life cycle of their host plant without causing any apparent pathogenesis [1]. Plant–microbe interaction studies have shown the contribution of microbial communities to the defence mechanisms of plants, as well as the substantial beneficial effects they can have on their host plant including improved acquisition of nutrients, accelerated growth, resilience against pathogens, and improved resistance against abiotic stress conditions such as heat, drought, and salinity [2].

The diversity and the structure of endophytic microbiome are dynamic and directly affected by ecological characteristics of the plant and soil such as the plant’s geographic location, the environmental factors and interactions within the host plant [3]. Specific members of the endophytic microbiome are part of a core microbiome [4] and most characterized members of endophytic microbial communities belong to the Actinobacteria, Bacteroidetes, Firmicutes, and Proteobacteria phyla [3, 5, 6]; however, endophytic microbiome structure can be affected by the host plant species, genotype (e.g. innate immunity receptors), plant organ or tissue type, developmental stage, growing season, geographic and field conditions, soil type, nutrient status of the host species and cultivation practices [2, 7, 8].

Endophytic microbes hold an enormous potential to increase host health. Endophytic bacteria can be used to overcome the effect of salinity stress, promote plant growth and to increase plant biomass and nutrient uptake; these approaches can provide a beneficial and environmentally friendly solutions for a sustainable global food security [9–11]. For successful use of endophytes, we need a deeper understanding of the endophytic community composition and the mechanisms that underlie the plant growth promotion, in order to successfully select the most efficient bacterial strains. Selected bacterial strains can be used in various combinations as synthetic communities in top-down approach to affect and study the endophytic microbiome.

Within endophytic bacterial communities, members show a strong influence on each other, which could include antagonistic, competitive, and mutualistic interactions [4]. This is the result of the nutritional competition, exchange, and even interdependence where metabolite exchange among microbes facilitates growth of some microbial species and in turn can influence microbiome composition of a given host species and their effect on the host, and therefore determine the final effect of plant microbiota-interactions in given conditions [2]. This is important when introducing new species or communities into an agricultural field or when trying to isolate the causative beneficial species in complex microbiomes. The host “genotype” can also have a dramatic impact on individual microbial species; individual cultivars can influence the structure of microbial communities and even the beneficial effects of endophytic bacteria [2, 12–14]. Thus, for the utilization of endophytic bacterial strains, an optimum approach is to isolate key bacterial strains from wild crop relatives (WCRs) [15].

Halophytes could be valuable potential sources of novel endophytic strains that can be used to overcome salinity stress, overcome plant diseases and augment plant growth [16–19]. Since, soil salinity not only disrupts the physiological and morphological plant processes but also increases susceptibility to pathogens – a major hindrance to crop yields [19] – the use of plant growth-promoting endophytes, isolated from halophyte WCRs for use in crops, can be a more eco-friendly approach than the use of agricultural chemicals.

In the present study, we test the hypothesis that the cultivated endophytic bacteria isolated from three halophytic plant species, two of *Brassicaceae* family (or Cruciferae): *Matthiola tricuspidata* and *Cakile maritima*, and one of *Apiaceae* family: *Crithmum maritimum*, have properties of salinity stress tolerance, plant growth promotion and phytopathogen growth inhibition and could have a predominant role in the future for a sustainable crop production of crops belonging mainly but not exclusively to those two plant families. In order to test this hypothesis, we isolated, cultured and identified 115 different bacterial strains and functionally characterized them using *in-vitro* and *in-planta* assays. This is the first study of bacterial endophytes obtained from *Matthiola tricuspidata*, *Crithmum maritimum* and *Cakile maritima*, which identifies their potential use as bacterial inoculants in commercial plants in order to overcome salinity stress and important plant diseases caused by the economically important pathogens *Verticillium dahliae*, *Ralstonia solanacearum* and *Clavibacter michiganensis* ssp. *michiganensis*.

## Material and Methods

### Site description and plant sample collection

Samples were collected during summer 2018 in three distinct sites in the Crete, Greece: site 1 (S1: N35°25’. E24°41’), site 2 (S2: N35°06, E25°48) and site 3 (S3: N35°00, E25°44) (**Suppl. Figure S1**). At S1, a natural beach area that favours salt-marsh vegetation, three *Matthiola tricuspidata* individuals were collected. At S2, a beach area located in the beach area of Pachia Ammos village, three *Crithmum maritimum* individuals were collected. At S3, a popular beach area located in the town of Ierapetra, three individuals of *Cakile maritima* were collected. Each sample was collected with sterile tools, placed in separate plastic bags to avoid cross contamination and immediately transported to the laboratory for processing.

### Plant surface sterilization endophytic cell isolation

Leaf and root materials from each species were cut and processed individually. Plant material was gently washed with sterile distilled water repeatedly to remove soil and dust particles. For surface sterilization, plant roots and leaves were placed into sterile Erlenmeyer flasks containing ethanol 75% v/v^-1^ for 60 sec with shaking and then in sterile Erlenmeyer flasks containing sodium hypochlorite solution 3 % w/v^-1^ (NaClO) for 10 min. The plant materials were then placed again in ethanol 75% v/v^-1^ for 60 sec. To remove the remaining NaClO, roots and leaves were rinsed 10 times with sterile distilled water (dH_2_O). The sterilization and transfer procedures were carried out in a type II laminar flow hood. About 100 μl of the last rinse (for each analysed sample) was evaluated for surface sterilization efficiency by plating on NA medium and monitored for microbial growth. Approximately 500 mg of leaves, stems and roots per each species were weighed and slashed to small parts for further processing. Only successfully sterilized root material was used for further analysis.

From each sample 0.5 - 1 gr of fresh biomass was cut into smaller pieces using a sterile scalpel and further grounded into a slurry with an autoclaved pestle and mortar. The slurry was transferred into sterile petri dishes and 30 ml of autoclaved dH_2_O was added. The petri dishes were sealed and placed onto a rotary shaker (150 rpm) at 25 °C for 2 h. After shaking, 100 ul of the material in triplicate were plated on Nutrient Agar (NA) plates using spread plate technique. Plates were incubated at 28 °C. Unique bacteria from each plate were chosen based on colony colour and morphology. Pure bacterial colonies were grown in Nutrient Broth (NB) and cells stocks were stored in 50% v/v glycerol at −80 °C. A total of 115 strains were isolated.

### Bacterial isolation and strains identification

A total of 115 bacterial isolates were isolated from the various plant tissue samples, based on morphological criteria of their forming colonies and their identities confirmed by the 16S rDNA sanger sequencing method.

To extract crude genomic DNA 1 ml of liquid bacterial culture in NB medium, was placed in liquid nitrogen for 15 sec. After incubation in room temperature, the lysate was centrifuged at 10,000 × g for 1 min. Two μl of the lysate were used to amplify the 16S rDNA genes using primers 27F: 5′-AGAGTTTGATCCTGGCTCAG-3′ [20] and 1492R: 5′-GTTTACCTTGTTACGACTT-3′ as the reverse primer [21]. PCR reactions of 20 μL were amplified in a BioRad T-100 Thermocycler with initial denaturation at 94 °C for 2 min, followed by 35 cycles of 5 sec at 94 °C, 30 sec annealing at 47 °C, 2 min primer extension at 72 °C, and a final extension at 72 °C for 5 min. Apart from the lysate, each tube contained, Bac-Free PCR Buffer, 250 nM of each primer, 0.2 mM of each deoxyribonucleotide triphosphate and 0.1 U BAC-Free HotStart Taq polymerase (Nippon Genetics, Europe). The PCR products were purified using Nucleo Spin Gel and PCR Clean up (Macherey-Nagel, Germany). The cleaned-up PCR products were sent Macrogen (Europe) for sequencing with primer 27F.

The resulting chromatograms were quality inspected using MEGA5 [22] and the start/end regions of low quality were manually trimmed off. Cleaned-up fasta files were aligned in SILVA [23]. The resulting nearly complete sequences of the 16S rDNA gene were queried against ezBioCloud [24] reference database for identification and documentation of the described bacterial strain with the closest sequence similarity.

### Bacterial salt tolerance assay

The salt tolerance of all bacterial isolates was estimated on the basis of the population density of these strains at different concentrations of NaCl [(ranging from 0.5% 5%, 10%, 15% and 17,5% (w/v)] in NA medium. Sterilized petri plates containing 25 mL NA medium with increasing NaCl concentrations were inoculated with 10 μl drops of freshly prepared NB of all strains and incubated at 28 °C. For each NaCl concentration, an *Escherichia coli* laboratory strain was inoculated as a negative control. After 24 h of incubation, the growth of the culture was estimated compared to *E. coli* growth.

### In-vitro growth inhibition of phytopathogens

Antibacterial activity of the bacterial isolates against the phytopathogenic bacteria *Ralstonia solanacearum* and *Clavibacter michiganensis* was evaluated by co-culturing each of the 115 bacterial isolates on NA plate lawn covered by *R. solanacearum* or *C. michiganensis*. The inhibition zone indicating bacterial growth inhibition was recorded as the antibacterial effect. Antifungal activity of the isolates against *Verticillium dahliae* was investigated. PDA medium was incubated with each bacterial isolate for 24h at 28 °C and then *V. dahliae* was inoculated at room temperature for 3-4 weeks. Fungal growth inhibition was determined by measuring the inhibition zone of *V. dahliae* hyphae on the media.

### In-vitro hemolysis screening assay

Many mammalian pathogenic bacteria are hemolytic in nature and their ability to lyse red blood cells is important for virulence in several of these species, however, not all. In order to assay the bacterial isolates for hemolysis activity, each isolate was grown on blood agar plates. The bacterial strains were inoculated with the spot test method and were incubated at room temperature for 48 hrs. The known non-mammalian-pathogenic species *Ensifer meliloti* was employed as a negative control.

### In-vitro growth inhibition of fungal human pathogen

Antifungal activity of specific isolates against anthropopathogenic fungus *Aspergillus fumigatus* was evaluated by co-culturing 11 bacterial isolates on NA plate lawn covered by *A. fumigatus* for 72 hours in room temperature under absence of light. The following strains were tested: CML04, CMR11, CMR22, CMR25, CrR12, CrR25, MTR12, MTR17a, MTR17b, MTR17c, MTR17d. Fungal growth inhibition was determined by the growth inhibition zone of the *A. fumigatus* hyphae on the media.

### In-planta salt tolerance assays

Twelve of the isolated bacterial strains were selected, according to their ability to grow in high salinity conditions (up to 17.5% w/v NaCl), in order to test their capacity to promote plant growth of the model plant *Arabidopsis thaliana*. Firstly, the experiment was performed with no abiotic stress conditions. The bacterial strains were cultured in Nutrient Broth medium for 46 h at 25 °C with stirring. The liquid cultures were centrifuged at 224 × g for 15 min, the supernatant was discarded and the cells were resuspended in 50 ml dH_2_O. Seeds of *Arabidopsis thaliana* ecotype Columbia (Col-0) were grown in plastic pots (6 × 6 × 7 cm) filled with vermiculite: soil (1:1), at 25 °C (16 h light / 8 h dark). For each bacterial strain and the corresponding control, 5 individual plants were grown in each pot. *A. thaliana* plants were watered with dH_2_O for 10 days. Then, plants were watered with 10 ml suspensions of the 12 re-dissolved bacterial strains for 7 days in order to let the bacterial strains to adapt. Following that, plants were watered with 10 ml dH_2_O every 2-3 days for 30 days. At the end of the treatment, the fresh weight of the leaves of each plant was measured. The leaves were then dried at 65-70 °C for 2 days and their dry weight was measured.

The same experiment was performed under salt treatment. Specifically, after the 7-day period of bacterial strain inoculation, instead of dH_2_O, the plants were watered with 10 ml of 200-250 mM NaCl. Fresh and dry weight of the leaves as measured.

For both experiments, mock samples were employed were no bacterial strains were inoculated and control plants that were inoculated with the strain *Escherichia coli* (Control-*E. coli*). The last control was used to check that the plants would not use the bacteria as a fertilizer.

### Confrontation and volatile tests of selected bacterial strains against V. dahliae

Direct *in vitro* antagonism of *V. dahliae* by selected bacterial strains was evaluated by dual-culture assays (confrontation test) on potato dextrose agar (PDA) (Lahlali et al., 2007). In particular, a 6-mm diameter mycelial disc taken from the periphery of a 2-week-old PDA culture of the fungus was placed in a new PDA plate (90 mm in diameter) at approximately 25-mm-distance from the center of the plate. Then, a 30-mm-long line from each bacterial strain (taken from a 48-h-old TSB liquid culture with an inoculation loop) was streaked on the opposite site of the plate at equal distance from the center (one strain per plate). Moreover, *Trichoderma harzianum* strain T22 was isolated from the commercial biofungicide TRIANUM-P (Koppert B.V. Hellas) and included in *in vitro* bioassays for comparison. Plates inoculated only with *V. dahliae* agar discs were served as controls. Plates (three per bacterial strain plus controls) were incubated at 24 °C in the dark. The radius of fungal colonies towards the direction of the test strain and that of controls was measured 5, 7, 9 and 12 days post inoculation (d.p.i.) and radial growth rates were expressed in mm/day. At the end of the bioassays (12 d.p.i.) the underside of the plates was scanned using a Samsung Xpress SL-M2875ND Laser Multifunction Printer at 1200 dpi and microsclerotial area on each plate image was determined manually using the image processing software ImageJ 1.46r (National Institutes of Health, USA). Then, the number of spores was estimated by transferring a 6-mm-diameter disc taken from the periphery of each culture into a 1.5 ml Eppendorf tube with 1 ml of water, and vortexed for 30 seconds. The number of spores was measured with the use of a haematocytometer under a light microscope. Moreover, actively growing mycelia from cultures’ periphery (located closer to test strain) were prepared and microscopic observations (30 readings per culture) were carried out to estimate hyphae width.

To evaluate the capacity of bacterial strains to affect *V. dahliae* growth via the production of volatile compounds, dual-plate assays [25] were conducted (volatile test). In brief, one 6-mm-diameter agar disc of actively growing mycelium of the fungus was placed in the center of a new PDA plate (90 mm in diameter), whilst each bacterial strain (taken from a 48-h-old TSB liquid culture) was streaked on another PDA plate. Then, the covers of the two plates were removed and resultant plates were adjusted together (bacterial culture was upturned) and sealed with cellophane membrane so the two microbes would share the same headspace without coming in contact with each other. Dual plates (upright and upturned) inoculated only with *V. dahliae* were served as controls. Likewise, in dual-culture assays, dual-plates (three per bacterial strain) were incubated at 24 °C in the dark and the radial growth, microclerotial area, sporulation and hyphae width of fungal colonies was measured as described above.

Radial growth inhibition (RGI), microsclerotia formation inhibition (MFI), sporulation inhibition (SI) and hyphae thinning (HT) were calculated according to the formula: ((Vc-Vt)/Vc)×100 where Vc= the microscopic value of *V. dahliae* in control plates and Vt= the respective value of *V. dahliae* against the antagonistic strain in dual-culture or dual-plate assays.

### Bacterial strains and fungal inoculum preparation for in-planta bioassays

Fifteen bacterial strains (assigned as CrR14, CrR18, CrR04, MTR12, MTR18, CMR01, CMR03, CML04, CMR25, MTR17a, MTR17d, MTR17f, MTR17g, MTR17h and MTR17b, MTR17c) were used in *in-planta* bioassays. The strains were grown in tripticasein soy broth (TSB) liquid culture in Erlenmeyer flasks of 500 ml capacity, containing 200 ml of the medium, in an orbital incubator at 180 rpm and 28 °C for 48 h in the dark. Bacterial suspensions were centrifuged at 3000 × g for 10 min and cells were re-suspended in water reaching a final concentration of 10^8^ cfu ml^-1^ (measured by dilution plating).

The highly virulent *V. dahliae* isolate 999-1 [26], which originated from a symptomatic eggplant (*Solanum melongena* L.), was used. *Verticillium dahliae* conidial suspension for eggplant—*V. dahliae* bioassays was prepared according to Markakis et al. (2016). In brief, conidia were produced by growing each *V. dahliae* strain in potato dextrose broth (PDB) at 160 rpm and 25 °C in the dark for 5 days. Then, conidia were harvested by filtration through three layers of cheesecloth and the suspensions centrifuged at 3000 × g for 10 min. Spores were re-suspended in sterilized distilled water and their concentration was adjusted to 5 × 10^6^ conidia ml^-1^.

### In planta verticillium wilt suppression bioassays

Eggplant seedlings (cv. Black Beauty) were used in the *in-planta* bioassays. Plants at the one-true-leaf stage, grown in 100 ml-capacity pots containing soil substrate (HuminSubstrat, Klasmann-3 Deilmann GmbH, Germany) were root-drenched with bacterial suspension (20 ml of 10^8^ cfu ml^-1^ of each strain per plant), whereas plants that served as controls (negative=no bacterium/no fungus assigned as ‘C-’ and positive=no bacterium/plus fungus assigned as ‘V.D.’) were treated with 20 ml of water. One week later, eggplants (at the second-true-leaf stage) were inoculated with *V. dahliae* by drenching the soil substrate in each pot with conidial suspension (20 ml of 5 × 10^6^ conidia ml^-1^ per pot). Plants that served as negative controls (C-) were treated with 20 ml of water. Eggplants were maintained under controlled conditions at 23 ± 2 °C with a 12-h light and dark cycle.

Two independent experiments (experiments I and II) were conducted to evaluate the suppressive effect of the aforementioned bacterial strains against *V. dahliae*. In experiment I, eleven treatments (C-, V.d., V.d.+CrR14, V.d.+CrR18, V.d.+CrR04, V.d.+MTR12, V.d.+MTR18, V.d.+CMR01, V.d.+CMR03, V.d.+CML04 and V.d.+CMR25) were applied; whereas in experiment II, ten treatments were conducted (C-, V.d., V.d.+MTR17a, V.d.+MTR17d, V.d.+BMTR17f, V.d.+MTR17g, V.d.+MTR17h, V.d.+MTR17b, V.d.+MTR17c and V.d.+TRIANUM-P). The commercial biofungicide TRIANUM-P (Koppert B.V. Hellas) was also included in experiment II (assigned as V.d.+TRIANUM-P) and applied at the appropriate dosage according to manufacturer’s instruction (20 ml of 3 × 10^7^ cfu ml^-1^ per plant). TRIANUM-P was served as a *V. dahliae*-suppressive reference treatment in the present study. Within each experiment, each treatment consisted of seven plants and the experiments were replicated three times (21 plants per treatment in total).

### Disease assessment

Verticillium wilt symptoms on eggplant were recorded at 2-, 3- and 4-day intervals from 12 to 30 days post inoculation with *V. dahliae* (d.p.i.). Bioassays were evaluated by estimating disease severity, disease incidence, mortality and relative area under disease progress curve (RAUDPC). Disease parameters were recorded according to Markakis et al. [56]. In brief, disease severity at each observation was calculated from the number of wilting leaves, as a percentage of total number of leaves of each plant. Disease ratings were plotted over time to generate disease progress curves. Subsequently the area under disease progress curve (AUDPC) was calculated by the trapezoidal integration method [27]. Disease was expressed as a percentage of the maximum possible area with reference to the maximum value potential reached over the whole period of each experiment and is referred to as relative AUDPC (RAUDPC). Disease incidence was estimated as the percentage of infected plants. Only plants with a final disease severity of ≥20% were considered infected, in order to discriminate between *V. dahliae*-associated disease symptoms and other weak symptoms occasionally observed (**Supplementary Table S2**). Mortality was estimated as the percentage of dead plants.

### Plant growth

Growth parameters were evaluated at the end of bioassays (at 24 and 30 d.p.i. for experiments I and II, respectively). To estimate the effect of the aforementioned treatments on plant growth, all plants were clipped off at the soil surface level and their height, fresh weight and number of leaves were measured.

### Fungal pathogen re-isolation

To verify the presence of the applied *V. dahliae* strain in plant tissues, five plants per treatment in each experiment were randomly selected. Specifically, the leaves of eggplants which had been cut above the soil level previously were removed and their stems were surface-disinfected by spraying with 95% ethyl alcohol and by quickly passing them through flame three times. For each plant, 3 xylem chips taken from different sites along the stem (base, middle and upper part of the stem) were aseptically placed onto acidified potato dextrose agar (PDA) after the removal of the phloem. Plates were then incubated at 24 °C in the dark for 14 days. The emerging fungi that grew out of tissue excisions were examined visually and under a light microscope and identified according to their morphological characteristics [28]. Pathogen isolation ratio was expressed as the frequency of positive *V. dahliae* isolations of each plant.

### Statistics

Analysis of variance (ANOVA) was employed to determine the effects of replication (1, 2 or 3), treatment (C-, V.d., V.d.+CrR14, V.d.+CrR18, V.d.+CrR04, V.d.+MTR12, V.d.+MTR18, V.d.+CMR01, V.d.+CMR03, V.d.+CML04, V.d.+CMR25 in Experiment I and C-, V.d., V.d.+MTR17a, V.d.+MTR17d, V.d.+BMTR17f, V.d.+MTR17g, V.d.+MTR17h, V.d.+MTR17b, V.d.+MTR17c, V.d.+TRIANUM-P in experiment II) and their interaction on disease incidence (DI), final disease severity (FDS), mortality (M), relative area under disease progress curve (RAUDPC) and isolation ratio (IR), and on plant height, fresh weight and total number of leaves (**Supplementary Tables S2 & S3**). Prior to ANOVA, homogeneity of variance across treatments was evaluated and an arcsine transformation was applied to normalize variance. When a significant *F* test was obtained for treatments (*P ≤* 0.05), the data were subjected to means separation by Tukey’s honestly significant difference test. Morphological and physiological characteristics of *V. dahliae* in dual-culture and dual-plate assays were also analysed by Tukey’s test (*P ≤ 0.05*). Moreover, standard errors of means were calculated.

### Bacterial total genome sequencing

Genome sequences of twelve bacterial isolates were determined using a 250 bp paired-end library with the Illumina MiSeq sequencing system (University of Exeter Sequencing Service, Exeter, UK). Reads were assembled using SPAdes 3.12.0 [29] and the assembled sequence annotated using the NCBI Prokaryotic Genome Annotation Pipeline (PGAP). Raw sequence reads and assembled genomes were uploaded to the Sequence Read Archive [30] and GenBank [31] and are available under BioProject accession number PRJNA634334.

### Bacterial genome analysis and annotation

For the bacterial genome analysis, we used the “RAST” (Rapid Annotation using Subsystem Technology) annotation server [32] a web service which can provide a quick and reliable genome annotation.

### Sequence alignment and phylogenetic tree construction

Alignments of the selected genes sequences were performed by ClustalX v2.0 [33] and followed by manual corrections. Sequence relationships were inferred using the maximum-likelihood method. Maximum-likelihood phylogenies were constructed using MEGA 5.2 [34]. genetic-trees were constructed assuming the bootstrap value derived from 1,500 to represent the evolutionary history of the included taxa. For the construction of genetic trees we used the concatenated *recA* and *gyrB* genes.

## Results

### Isolation and grouping of culturable bacteria

After surface sterilization, no epiphytic bacteria could be cultivated from the surface of the roots and leaves of halophytic plants. Endophytic bacteria were cultivated from all three halophytes and in total, 115 pure cultures showing different colony morphology per isolation source (plant, root or leaf) were obtained; 91 were retrieved from roots and 24 from leaves. From *M. tricuspidata* 45 strains were isolated, from *C. maritima* 31 strains, and from *C. maritimum* 39 strains (**Supplementary Table S1).**

For the 115 isolates, the sequencing of 16S rRNA gene amplicons, yielded sequences of sufficient length for taxonomic analysis (**Figure 1, Supplementary Table S1**). The distribution and abundance of bacteria varied according to the type of samples (**Figure 1**). The bacterial isolates were assigned to five different classes (**Supplementary Table S1**) and 24 genera (**Figure 1, Supplementary Table S1**). The most prevalent genus among the isolated bacteria was *Bacillus*, accounting for 24% of total strains, followed by *Enterobacter* (19%), *Pseudomonas* (12%), *Microbacterium* (6%), *Paenarthrobacter* (6%), and *Pantoea* (5%). The highest number of isolated bacteria were hosted in roots of *M. tricuspidata* (38), followed by *C. maritima* roots (28) and *C. maritimum* roots (25) in contrast to the number of isolated bacteria from leaf samples (*M. tricuspidata*: 7, *C. maritima*: 4 and *C. maritimum*: 13). *Bacillus* was isolated from all plants and tissues except the roots of *C. maritimum. Enterobacter* and *Pseudomonas* strains were only isolated from root samples.

**Figure 1.**
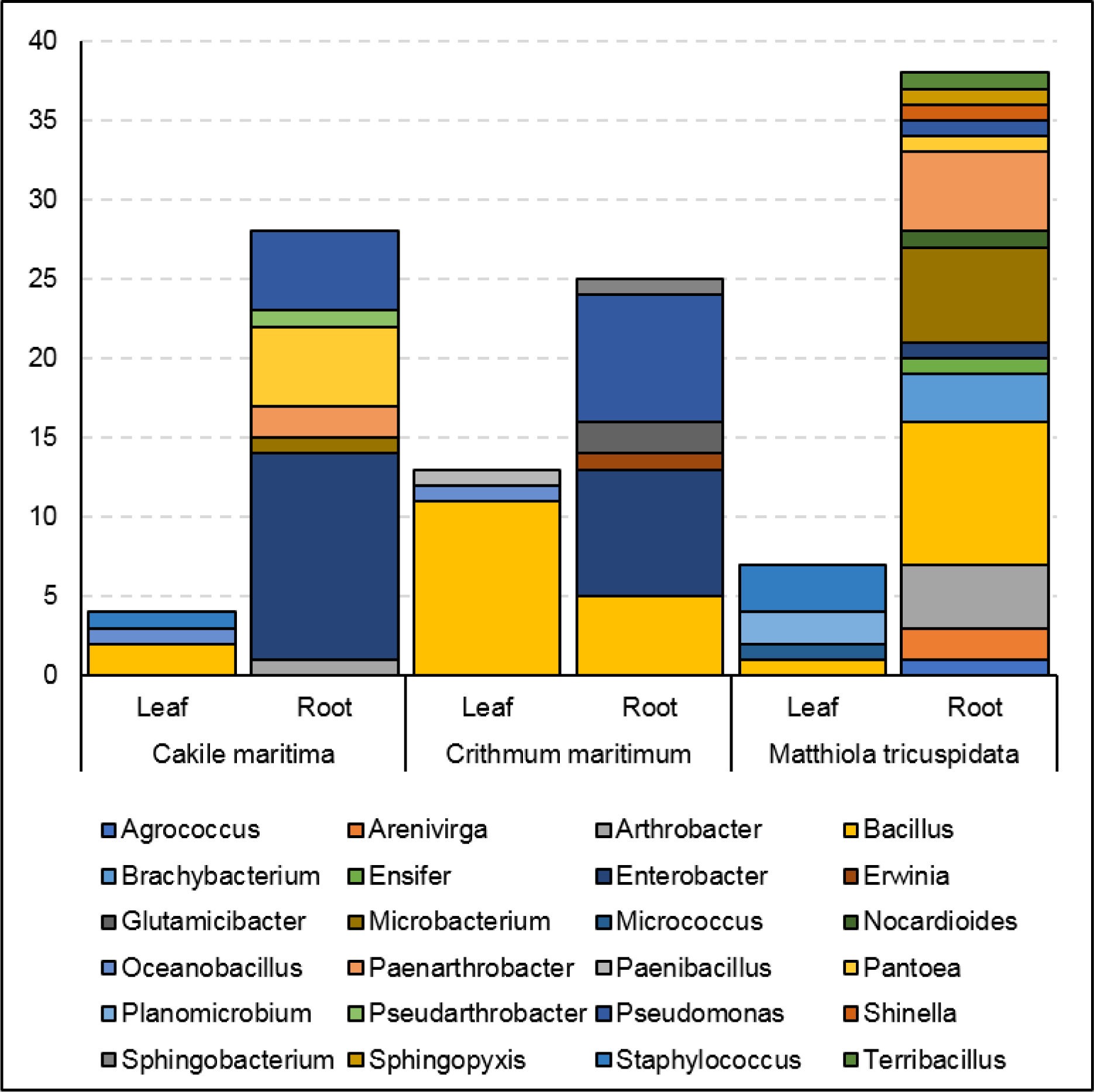
Abundance of genera of bacterial isolates obtained from leaves and roots of *tricuspidata, Crithmum maritimum* and *Cakile maritima* plants.

### Shared and unique endophyte taxa among plants and plant compartments

Except *Bacillus* isolates, which were isolated both from roots and leaves, the rest of the genera with more than one isolate were only found in either roots or leaves (**Supplementary Table S1**). Specifically, the second-most abundantly isolated genus, *Enterobacter,* was isolated only from root samples (**Supplementary Table S1**). Similarly, all strains from the genera *Enterobacter, Pseudomonas, Microbacterium, Paenarthrobacter, Pantoea, Arthrobacter, Brachybacterium, Arenivirga* and *Glutamicibacter* were isolated from root samples, whereas the strains *Oceanobacillus, Planomicrobium* and *Staphylococcus* were isolated only from leaf tissue samples (**Supplementary Table S1**).

The aforementioned most abundant culturable genera were hosted in at least two of the three halophytic species. *M. tricuspidata* hosted the biggest number of culturable genera (18), whereas *C. maritima* and *C. maritimum* hosted 10 and 8 different genera, respectively. *C. maritima* hosted the biggest number of *Enterobacter* isolates (13 of the total 22) and the smallest number of *Bacillus* isolates (2 out 28). Strains in the genera *Brachybacterium* and *Arenivirga* were isolated only from *M. tricuspidata* plants, whilst the *Glutamicibacter* isolates were hosted only in *C. maritimum* and the *Planomicrobium* isolates in *C. maritima*.

### Functional characteristics of isolates

Bacterial features known to contribute to stress tolerance or biocontrol were tested for their ability to grow in increasing salinity levels (5%, 7.5%, 10%, 15%, 17.5%) and their phytopathogen antagonistic activity against known phytopathogenic bacteria and fungi. The results are summarized in **Supplementary Table S1.**

The most common trait was growth at the salinity level of 5% (96 isolates), followed by the ability to inhibit the growth of the phytopathogenic fungi *Verticillium dahliae*, the ability to grow at 7.5% and 10% salinity, the ability to inhibit the phytopathogenic bacterium *Ralstonia solanacearum* and the ability to inhibit the growth of the bacterial phytopathogen *Clavibacter michiganensis* subsp. *michiganensis* (**Supplementary Table S1**).

Most of the bacterial isolates managed to grow at 5% salinity (96 strains). From the 28 strains isolated from the roots of *C. maritima*, 26 strains (92.9% of the total) showed ability to grow at 5% salinity, and 14 of these (50%) managed to grow at 7.5% salinity; the 4 strains isolated from the leaves of the same plant all 4 managed to grow at 10% salinity and 2 of these could grow at 17.5 %. 21 out of 25 strains (84%) of the roots of *C. maritimum* could grow at 5% salt and 16 (64%) could grow at 10% salinity. 12 out of 13 (92.3%) strains from the leaves of *C. maritimum* could grow at the 5% level and 8 (61.5%) could grow at 10% salt. From the 38 strains isolated from the root of *M. triscupidata*, 27 (71%) could grow at the 5% salt threshold and 3 could grow in 10% salt.

Of the 6 strains that managed to grow at the 17.5% salinity threshold, 4 were isolated from leaf tissues (**Supplementary Table S1**): *Staphylococcus saprophyticus* (CML12) and *Oceanobacillus picturae* (CML15) isolated from *C. maritima* leaves, *Oceanobacillus picturae* (CrL11) from *C. maritimum* leaves and *Micrococcus aloeverae* (MTL04) from *M. triscupidata* leaves (**Supplementary Table S1**). The two strains from root tissues that could grow on 17.5% are *Enterobacter hormaechei* subsp. *hoffmannii* (CMR13) isolated from *C. maritima* and *Bacillus hwajinpoensis* (CrR23) isolated from *C. maritimum* (**Supplementary Table S1**). Another 3 additional Bacilli strains (CrR16: *Bacillus haikouensis*, CrR22: *Bacillus haikouensis*, MTR05: *Terribacillus saccharophilus*) that could grow on 15% salinity, did not grow on 17,5% salinity (**Supplementary Table S1**).

### Phytopathogens growth inhibition ability

All 115 bacterial strains were subjected to *in vitro* inhibition assays against 3 known phytopathogens: the bacteria *Ralstonia solanacearum* and *Clavibacter michiganensis* subsp. *michiganensis* and the fungus *Verticillium dahliae*. In the assays against the phytopathogenic bacteria, the bacterial isolates that showed any kind of inhibition were characterized as having “weak”, “medium” or “strong” inhibitory activity based on the size of the inhibition zone around the bacterial colony (**Figure 2**). In the *in-vitro* assay against *Verticillium dahliae* the inhibitory activity was similarly judged as “weak”, “medium” or “strong” based on the linear distance between the bacterial and the fungal colonies (Figure 2).

**Figure 2.**
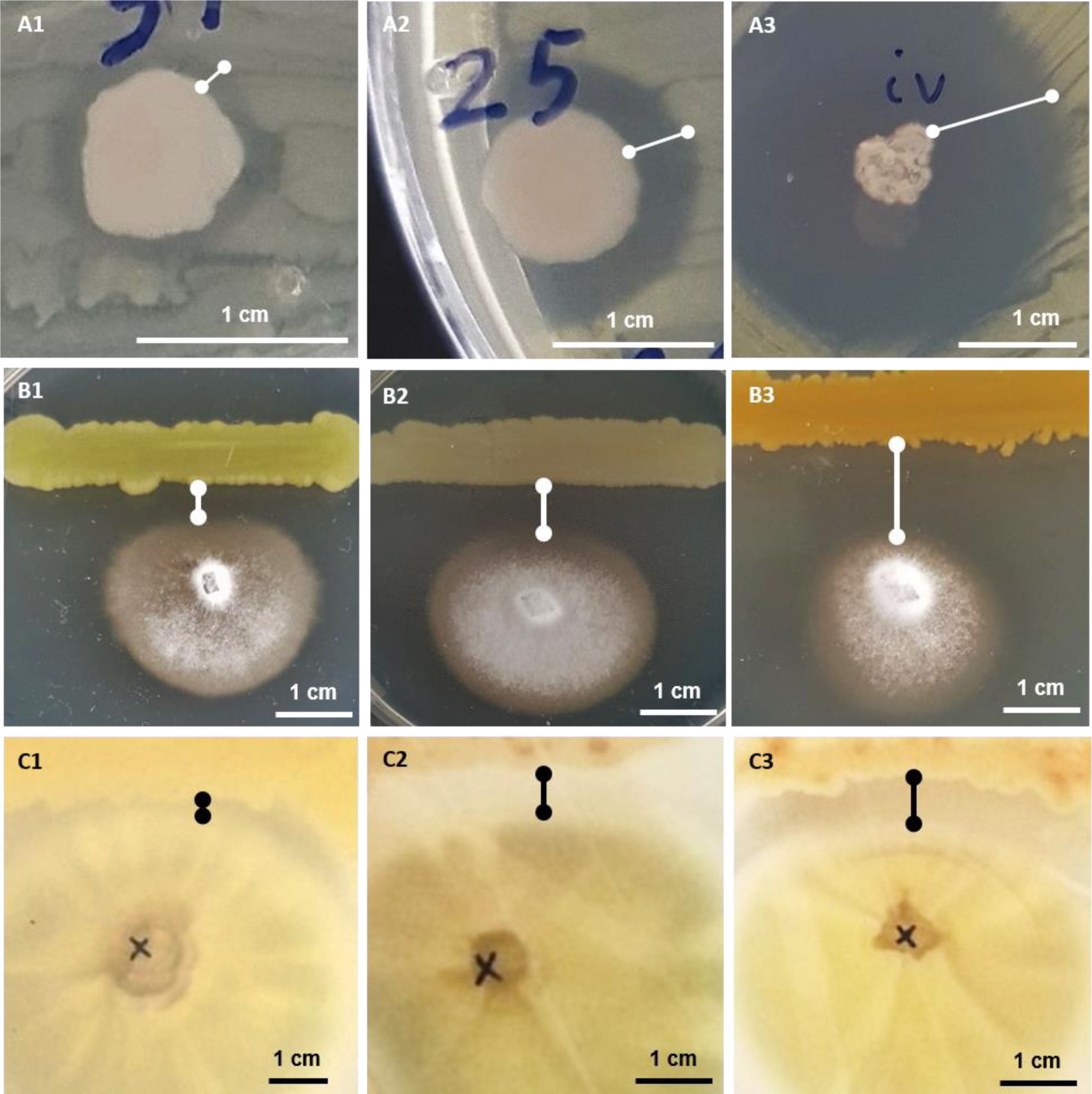
*In vitro* growth inhibition experiments: against phytopathogenic bacteria *Ralstonia solanacearum* or *Clavibacter michiganensis* (**A**), against phytopathogenic fungus *Verticillium dahliae* (**B**) and against human pathogenic fungus *Aspergillus fumigatus* (**C**). Representative experiments are shown to display their “small” (**A1, B1, C1**), “medium” (**A2, B2, C2**) or “large” (**A3, B3, C3**) inhibitory activity.

**Figure 3.**
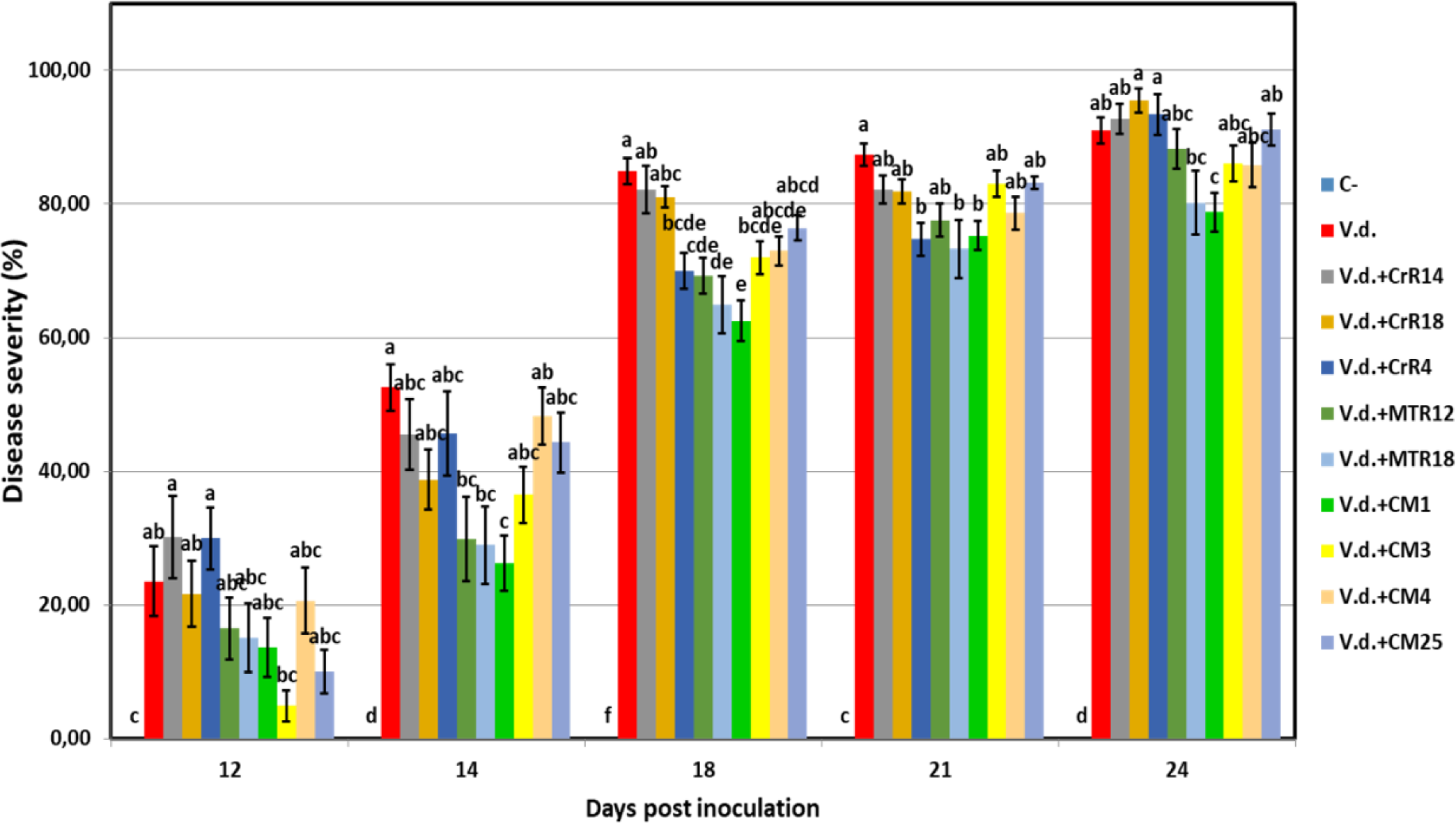
Verticillium wilt disease severity index on eggplant treated with various bacterial strains at 12, 14, 18, 21 and 24 days post inoculation with *Verticillium dahliae* conidial suspension (20 ml of 5 × 10^6^ conidia ml^-1^). Each column represents the mean of 21 plants after combining the results of 3 replicated experiments (experiment I). Columns at each observation time point followed by the same letter are not significantly different according to Tukey’s HSD test at *P ≤* 0.05. Vertical bars indicate standard errors.

Only 25 (21.7%) of the 115 bacterial strains demonstrated an inhibition zone of the *Ralstonia solanacearum* growth (**Supplementary Table S1**). These 25 strains belong to six genera: *Bacillus, Enterobacter, Erwinia, Glutamicibacter, Paenarthrobacter* and *Pseudomonas*. Only one strain isolated from leaf tissues of the 3 halophytes showed any kind of inhibitory activity; *Bacillus altitudinis* (strain CML04) isolated from *C. maritima*. Of the 45 strains isolated from *M. triscupidata* plants, 3 strains showed a weak inhibitory zone against *Ralstonia*. A total 10 strains (7 from *C. maritima* and 3 from *C. maritimum*), that belong to either to the genera *Enterobacter* and *Pseudomonas* (**Supplementary Table S1**), showed a strong inhibition zone (**Figure 2**).

An even lower number of strains showed any kind of inhibition against the phytopathogenic *Clavibacter michiganensis* subsp. *michiganensis*: a total 17 strains out of 115 (14.8%). All 9 isolates from *M. triscupidata*, that are *Bacillus licheniformis* or *Bacillus sonorensis* strains, demonstrated a strong inhibition zone (**Figure 2**). Similarly, 2 strains from *C. maritimum* roots (both *Pseudomonas glareae*, CrR12 and CrR13) showed a strong inhibition zone (**Supplementary Table S1**). An additional 2 and 4 strains showed weak and medium inhibition zone against *Clavibacter*, respectively (**Supplementary Table S1**).

Interestingly, a large majority (76.5%) of the bacterial isolates demonstrated some kind of inhibition of the phytopathogenic fungus *Verticillium dahliae* (**Supplementary Table S1**). These strains originate from both leaf and root tissue from all 3 halophytes. A strong inhibition zone was demonstrated by 34 of these 88 strains, all of which except one, were isolated from the roots of halophytic plants (**Supplementary Table S1**). From these 34, 11 strains belong to the genus *Bacillus*, another 11 to *Pseudomonas*, 5 to *Enterobacter*, 2 to *Microbacterium* and the rest belong *Agrococcus*, *Arthrobacter*, *Paenarthrobacter*, *Sphingobacterium* and *Sphingopyxis* (**Supplementary Table S1**).

Eleven of the strains were tested for inhibition against the human pathogenic fungus *Aspergillus fumigatus*. Interestingly, five of these strains were able to inhibit the growth of *A. fumigatus* (**Figure 2**). The strain CML04 (*Bacillus altitudinis*) isolated from *C. maritima* was the only strain isolated from leaf tissues that was able to show inhibitory effect, while all rest strains were isolated from the roots of *M. triscupidata* (**Supplementary Table S1**). Two strains MTR17d (*Bacillus sonorensis*) and MTR17b (*Bacillus licheniformis*) showed a strong zone of inhibition (**Figure 2, Supplementary Table S1**).

### In planta assay for plant growth promotion and salt tolerance

Bacterial isolates with salt tolerance between 10% and 17.5% NaCl concentration at the *in-vitro* assays were selected for the *in-planta* assays regarding their plant growth promotion under “no stress”. The model plant species *Arabidopsis thaliana* (*Brassicaceae*) was imbued with bacterial cultures for 7 days. After this period where the bacteria were left to adapt, the plants were watered for a month where the fresh and dry weight of the leaves was calculated (**Table 1**). As negative controls, dH_2_O, as well as, *E. coli* cultures were used.

**Table 1.** In planta plant growth promotion and salt tolerance assays in *Arabidopsis thaliana* plants. Plants were watered with selected halotolerant strains and with and without NaCl solution every 2-3 days for 30 days and fresh and dry leaf weight was measured.

| Host Plant | Strain | Plant growth promotion assay |  | Salinity stress assay |  |
| --- | --- | --- | --- | --- | --- |
|  |  | Fresh weight | Dry weight | Fresh weight | Dry weight |
| *Cakile maritima* | CML12 | 0.410 | 0.029 | 0.246 | 0.034 |
| *Cakile maritima* | CML15 | 0.204 | 0.025 | 0.309 | 0.052 |
| *Cakile maritima* | CMR13 | 0.144 | 0.022 | 0.239 | 0.032 |
| *Crithmum maritimum* | CrL01 | 0.541 | 0.053 | 0.147 | 0.031 |
| *Crithmum maritimum* | CrL04 | 0.149 | 0.019 | 0.197 | 0.031 |
| *Crithmum maritimum* | CrL11 | 0.242 | 0.026 | 0.236 | 0.034 |
| *Crithmum maritimum* | CrR16 | 0.277 | 0.026 | 0.165 | 0.031 |
| *Crithmum maritimum* | CrR22 | 0.094 | 0.016 | 0.275 | 0.042 |
| *Crithmum maritimum* | CrR23 | 0.170 | 0.023 | 0.103 | 0.026 |
| *Matthiola tricuspidata* | MTL01 | 0.113 | 0.017 | 0.202 | 0.037 |
| *Matthiola tricuspidata* | MTR05 | 0.169 | 0.028 | 0.322 | 0.056 |
| *Matthiola tricuspidata* | MTR27 | 0.114 | 0.015 | 0.213 | 0.033 |
|  | Control *E. coli* | 0.110 | 0.017 | 0.235 | 0.034 |
|  | Control H<sub>2</sub>O | 0.156 | 0.023 | 1.266 | 0.094 |
|  | Control Salt | N/A | N/A | 0.25 | 0.034 |

The same experiment was repeated where after the 7-day mark, the plants were watered with 10 ml of 250 mM NaCl solution every 2-3 days for 30 days. Except the aforementioned controls, an additional control of salty water was implemented.

Under no stress conditions, plants inoculated with strains CML12, CML15, CrL01, CrL11, CrR16, CrR23, MTR05 showed an increase in fresh leaf weight between 1.1 and 2.6 times to the non-inoculated plants and between 1.0 and 2.3 times increase in dry leaf weight (**Table 1**).

Under salt stress the growth promotion effect is less accentuated, since the plants that were imbued with strains CML15, CrR22, MTR05 had increased fresh leaf weight 1.2, 1.1, 1.3 times, respectively, in comparison to the salt control and 1.5, 1.2, 1.6 times increase in dry leaf weight, respectively (**Table 1**).

Strains CML15 and MTR05 conferred an increase in fresh and dry leaf weight both under no stress and under salt stress whereas strain CrR22 had a positive affect only under salt stress condition (**Table 1**). On the other hand, strains CML12, CrL01, CrL11, CrR16 and CrR23 had a positive effect on fresh and dry weight under no stress condition (**Table 1**).

### Direct and indirect in-vitro effects on Verticillium dahliae growth

All bacterial strains with the exception of MTR17h inhibited significantly *V. dahliae* growth rate in dual-culture assays, however, only MTR17h, MTR17b and MTR17c could suppress fungal growth by means of volatile compounds. Likewise, nearly all strains were capable of inhibiting fungal sporulation (except of MTR17g) in dual-culture assays. While, most of them inhibited significantly spores production in dual-plate assays. Interestingly, some strains caused a significant induction of *V. dahliae* sporulation in such assays (MTR17h, MTR17b and MTR17c) indicating that fungal growth suppression induces fungal sporulation. Moreover, 6 out of 16 strains could reduce significantly hyphae width in direct culture conditions whereas 7 out of 16 were capable of reducing hyphae width by the mean of volatiles. In addition, 9 strains could inhibit microsclerotia formation significantly in dual-culture assays; however only 3 strains reduced microsclerotia formation significantly in dual-plate assays. Interestingly, MTR17h caused a significant induction in microsclerotia formation either in dual-culture or in dual-plate assays.

### Suppression of Verticillium wilt symptoms in-planta

For the suppression of *Verticillium dahliae* wilt symptoms *in-planta* we used a well-established fungus/plant system, the *Verticillium*/eggplant system. We selected a small number of our bacterial isolates that initially revealed a noteworthy *in-vitro* growth inhibition effect to *Verticillium*.

We performed two distinct assays that we mark hereafter as “experiment I” and “experiment II”. *V. dahliae* wilt symptoms on eggplant started 12 days after inoculation (d.p.i.), with *V. dahliae* conidial suspension and were recorded periodically for another 12 days in experiment I. Strains CrR4, MTR12, MTR18 and CMR01 suppressed significantly disease severity at 18 and 21 d.p.i. whereas MTR18 and CM1 treatments caused significant reduction of disease severity at most observation time points (**Table**). Taken all disease parameters into account, CMR01 was the most effective strain in terms of disease suppression (**Table 2**; **Supplementary Tables S2 & S3**). Likewise, first disease symptoms in experiment II were also observed on 12 d.p.i. and recorded until 30 d.p.i. Disease severity progressed rapidly in the control (*V.d.*) and the non-suppressive treatments (MTR17d, MTR17f and MTR17g), whereas MTR17a-, MTR17h-, MTR17b- and MTR17c-treated plants showed less prominent symptoms and slower disease development (**Table 2;** **Figure 5**). Data on various disease parameters indicated that strain MTR17h, is comparable to the positive control used in these assays, the fungus *Trichoderma harzianum* strain TRIANUM-P, both were the most effective in symptom suppression (**Table**). Interestingly, the observed decrease in symptom severity in MTR17h-treated plants was associated with significantly lower *V. dahliae* re-isolation ratio compared to positive control (*V.d.*) plants, indicating less active growth of the pathogen into the xylem vessels. The interesting observation here is that the MTR17h isolate does not reveal strong growth inhibition effect on *V. dahliae* in our in-vitro assays. This result could suggest the plant innate immunity activation/reinforcement effect by the particular bacterial isolate, which needs to be further investigated in the future. Neither symptoms nor positive isolations were observed in negative control plants.

**Figure 4.**
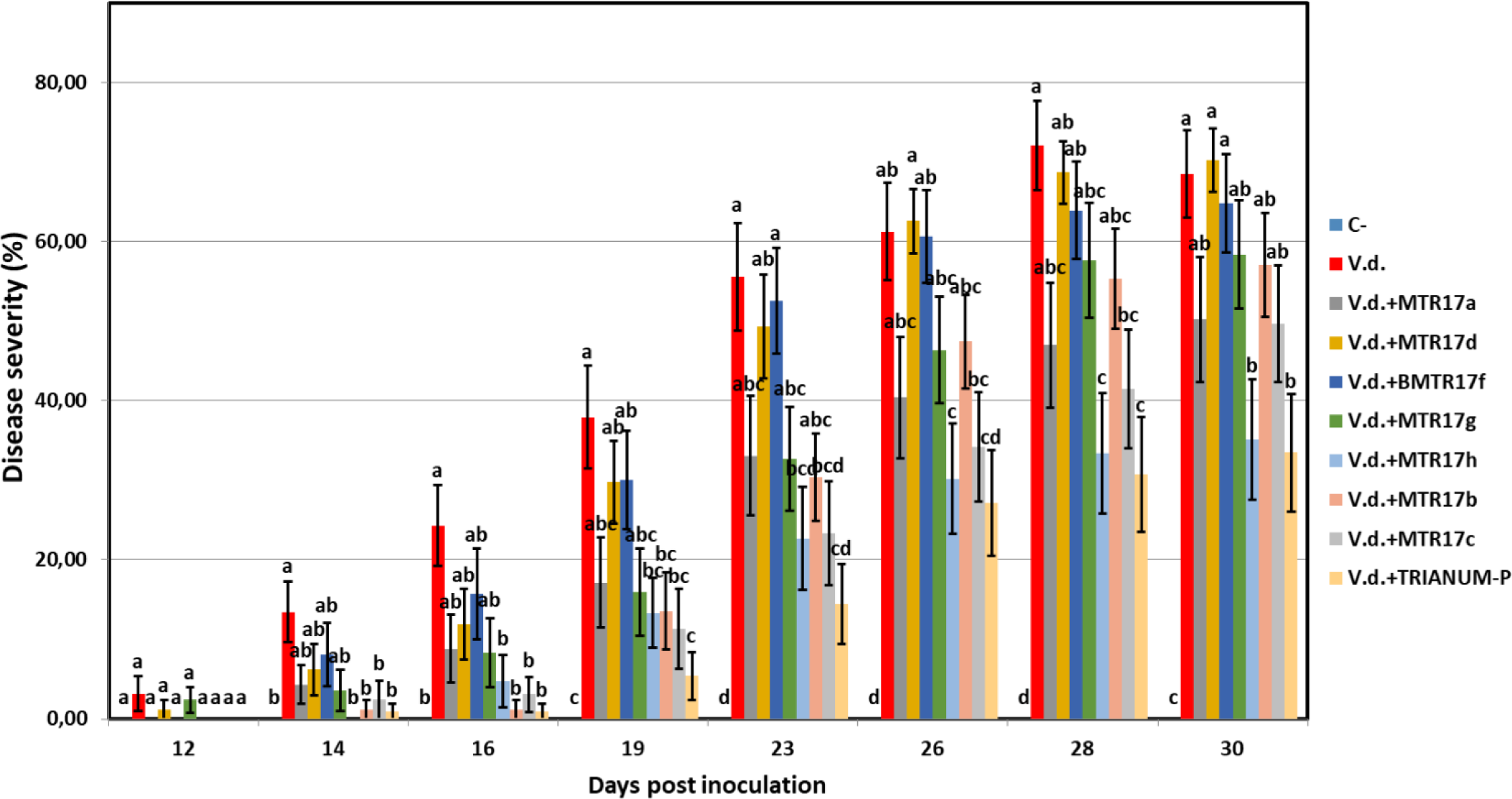
Verticillium wilt disease severity index on eggplant treated with various bacterial strains and the commercial biofungicide TRIANUM-P (Koppert B.V. Hellas) at 12, 14, 16, 19,23, 26, 28 and 30 days post inoculation with Verticillium dahliae conidial suspension (20 ml of 5 × 106 conidia ml-1). Each column represents the mean of 21 plants after combining the results of 3 replicated experiments (experiment II). Columns at each observation time point followed by the same letter are not significantly different according to Tukey’s HSD test at P ≤ 0.05. Vertical bars indicate standard errors.

**Figure 5.**
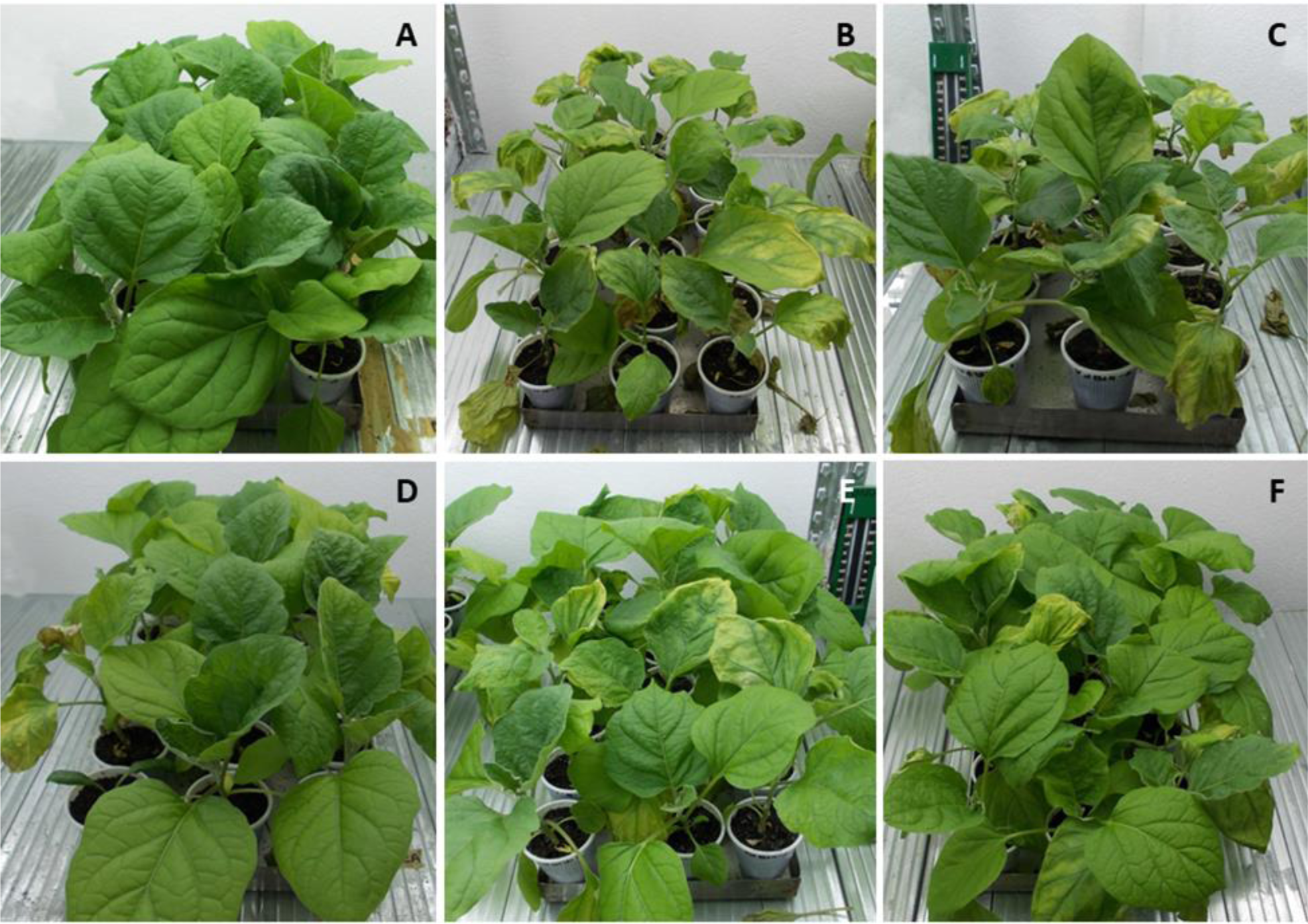
Verticillium wilt symptoms on eggplants treated with various bacterial strains and the commercial biofungicide TRIANUM-P (Koppert B.V. Hellas). **A:** mock inoculated plants treated with water (negative controls); **B:** *Verticillium dahliae*-inoculated plants with no other treatment (positive controls), **C:** *V. dahliae*-inoculated plants treated with the non-suppressive bacterial strain Mtr17d; **D and E:** *V. dahliae*-inoculated plants treated with the disease-suppressive strains Mtr17h and Mtr17c, respectively; **F:** *V. dahliae*-inoculated plants treated with the disease-suppressive biofungicide TRIANUM-P.

**Table 2.** Values of fungal parameters of *Verticillium dahliae* treated with 16 different bacterial strains (CrR14, CrR18, CrR4, MTR12, MTR18, CM1, CM3, CM4, CM25, MTR17a, MTR17d, MTR17f, MTR17g, MTR17h, MTR17b, MTR17c) and *Trichoderma harzianum* strain T22 in dual-culture and dual-plate assays. Values were estimated as the percentage of inhibition compared to control (*V.d.*)

| Treatment | Fungal parameters<sup>a</sup> |  |  |  |  |  |  |  |
| --- | --- | --- | --- | --- | --- | --- | --- | --- |
|  | Dual-culture assays (confrontation test) |  |  |  | Dual-plate assays (volatile test) |  |  |  |
|  | RGI (%) | SI (%) | HWT (%)<sup>b</sup> | MFI (%) | RGI (%)<sup>c</sup> | SI (%)<sup>d</sup> | HWT (%)<sup>e</sup> | MFI (%)<sup>f</sup> |
| V.d. | 0.00 h | 0.00 e | 0.00 d | 0.00 f | 0.00 cde | 0.00 b | 0.00 de | 0.00 cde |
| V.d.+CrR14 | 52.92 c | 70.87 cd | 25.21 abc | 68.98 ab | 8.75 c | 83.08 a | 18.46 abcd | 13.56 bcd |
| V.d.+CrR18 | 76.78 b | 57.11 d | 33.53 a | 52.41 bcde | -1.97 cde | 79.68 a | 22.33 ab | 24.99 cde |
| V.d.+CrR4 | 45.73cd | 74.30 bc | 20.56 abcd | 47.88 bcde | 4.66 cd | 21.84 ab | 21.09 abc | 71.91 ab |
| V.d.+MTR12 | 23.97 ef | 92.94 a | 22.92 abc | 26.65 cdef | -3.83 cde | 82.75 a | 25.99 a | -20.10 cde |
| V.d.+MTR18 | 33.95 de | 89.36 bc | 23.49 abc | 34.23 bcdef | -2.78 cde | 83.13 a | 30.95 a | -1.86 cde |
| V.d.+CM1 | 21.91 ef | 81.46 abc | 30.26 ab | 60.41 abcd | -1.54 cde | 84.81 a | 30.72 a | -25.08 cde |
| V.d.+CM3 | 27.22 ef | 88.34 abc | 17.56 abcd | 47.42 bcde | 7.70 c | 56.63 ab | 21.45 abc | 42.98 abc |
| V.d.+CM4 | 59.69 c | 70.31 cd | 18.10 abcd | 65.11 abc | -1.10 cde | 78.68 a | 28.22 a | -32.44 de |
| V.d.+CM25 | 23.15 ef | 79.00 abc | 13.67 abcd | 23.94 def | -3.58 cde | 70.63 ab | 17.55 abcd | -12.64 cde |
| V.d.+MTR17a | 45.22 cd | 82.66 abc | 6.89 cd | 22.63 def | -8.86 de | -100.42 c | -5.11 e | -43.35 cd |
| V.d.+MTR17d | 45.85 cd | 83.99 abc | 5.56 ab | 32.93 bcdef | 3.31 cd | 74.61 ab | -3.78 e | -2.18 de |
| V.d.+MTR17f | 51.46 c | 74.21 bc | 10.89 bcd | 18.77 ab | -16.21 e | 77.33 ab | 1.56 de | -60.99 e |
| V.d.+MTR17g | 16.75 fg | 5.29 e | 21.33 abc | 62.41 abcd | -4.50 cde | 38.33 ab | 3.00 cde | -2.99 cde |
| V.d.+MTR17h | 2.86 gh | 70.53 cd | 5.59 cd | -67.79 g | 55.99 b | -116.97 c | 5.59 bcde | -298.98 f |
| V.d.+MTR17b | 49.02 cd | 81.79 abc | 0.58 d | 22.91 def | 74.57 a | -120.55 c | 0.58 de | 97.83 a |
| V.d.+MTR17c | 60.21 c | 91.26 a | 7.60 cd | 24.02 def | 69.06 ab | -88.87 c | 6.35 bcde | 99.28 a |
| V.d.+TRIANUM-P | 95.79 a | 79.46 abc | nm | 98.91 a | ne | ne | ne | ne |

<sup>a</sup> Fungal parameters were calculated according to the formula: $((V_c - V_t)/V_c) \times 100$ where $V_c$ = the microscopic value of *V. dahliae* in control and $V_t$ = the respective value of *V.* *dahliae* towards the antagonistic strain in dual-culture or dual-plate assays. Each value represents the mean of 3 replicates. RGI=radial growth inhibition; SI=sporulation
(spore production) inhibition; HWT=hyphae width thinning; MFI=microsclerotia formation inhibition. Within columns, values followed by the same letter are not significantly different according to Tukey's HSD test at $P \leq 0.05$
<sup>b</sup> 'nm' indicates that HWT values were not measured since *T. harzianum* overgrown *V. dahliae* in dual-culture assays and pathogen hyphae could not be identified
<sup>c, d, e, f</sup> 'ne' indicates that RGI, SI, HWT and MFI were not estimated since *T. harzianum* could reach directly *V. dahliae* even in dual-plate assays

### Effects of treatments in plant growth

The measurements of growth parameters of eggplant inoculated with *V. dahliae* and treated with various bacterial strains and the *T. harzianum* strain TRIANUM-P or not (C-), are shown on **Table 3**. *Verticillium dahliae*-inoculated plants treated with MTR17c and *T. harzianum* TRIANUM-P developed significantly higher fresh weight compared with the *V. dahliae*-inoculated controls, whereas most of the plant growth parameters in non-inoculated plants were significantly higher than in the inoculated ones.

**Table 3.** Values (±standard errors) of disease parameters for eggplants inoculated with *V. dahliae* and treated with different bacterial strains and TRIANUM-P (CrR14, CrR18, CrR4, MTR12, MTR18, CM1, CM3, CM4, CM25 in experiment I, and MTR17a, MTR17d, MTR17f, MTR17g, MTR17h, MTR17b, MTR17c, TRIANUM-P in experiment II) or not (C-, V.d.)

| Experiment | Treatment | Disease parameters<sup>a</sup> |  |  |  |  |
| --- | --- | --- | --- | --- | --- | --- |
|  |  | DI (%) | FDS (%) | M (%) | RAUDPC (%) | IR |
| Experiment I | C- | 0.00±0.00 b | 0.00±0.00 c | 0.00±0.00 c | 0.00±0.00 c | 0.00±0.00 b |
| | V.d. | 100.00±0.00 a | 91.00±1.91 ab | 100.00±0.00 a | 42.36±2.08 a | 0.55±0.07 ab |
| | V.d.+CrR14 | 100.00±0.00 a | 92.74±2.21 ab | 95.24±4.76 a | 42.54±2.62 a | 0.65±0.05 a |
| | V.d.+CrR18 | 100.00±0.00 a | 95.50±1.80 a | 90.48±6.15 ab | 39.19±1.99 ab | 0.55±0.08 ab |
| | V.d.+CrR4 | 100.00±0.00 a | 93.40±2.99 a | 78.57±7.70 ab | 39.84±2.18 ab | 0.60±0.05 ab |
| | V.d.+MTR12 | 100.00±0.00 a | 88.29±2.74 abc | 80.95±9.91 ab | 33.86±2.59 ab | 0.80±0.07 a |
| | V.d.+MTR18 | 100.00±0.00 a | 80.19±4.75 bc | 71.43±8.69 ab | 31.67±3.02 b | 0.55±0.05 ab |
| | V.d.+CM1 | 100.00±0.00 a | 78.79±2.85 c | 52.38±14.29 b | 30.72±2.18 b | 0.55±0.08 ab |
| | V.d.+CM3 | 100.00±0.00 a | 86.07±2.71 abc | 66.67±14.55 ab | 32.25±1.47 b | 0.65±0.08 a |
| | V.d.+CM4 | 100.00±0.00 a | 85.82±3.34 abc | 69.05±5.67 ab | 37.91±2.00 ab | 0.65±0.05 a |
| | V.d.+CM25 | 100.00±0.00 a | 91.17±2.37 ab | 80.95±9.91 ab | 35.71±1.46 ab | 0.85±0.03 a |
| Experiment II | C- | 0.00±0.00 c | 0.00±0.00 c | 0.00±0.00 b | 0.00±0.00 d | 0.00±0.00 b |
| | V.d. | 90.48±6.15 ab | 68.49±5.43 a | 47.62±9.91 a | 26.76±2.95 a | 0.53±0.05 a |
| | V.d.+MTR17a | 71.43±15.31 ab | 50.21±7.88 ab | 33.33±10.29 ab | 15.05±2.81 bc | 0.20±0.09 ab |
| | V.d.+MTR17d | 95.24±4.76 a | 70.22±3.93 a | 23.81±14.02 ab | 23.04±2.20 ab | 0.40±0.20 ab |
| | V.d.+MTR17f | 85.71±9.91 ab | 64.79±6.12 a | 33.33±14.51 ab | 22.95±2.67 ab | 0.38±0.17 ab |
| | V.d.+MTR17g | 80.95±6.73 ab | 58.36±6.85 ab | 38.10±11.34 ab | 16.81±2.84 abc | 0.27±0.12 ab |
| | V.d.+MTR17h | 52.38±12.30 b | 35.15±7.58 b | 4.76±4.76 b | 10.51±2.58 cd | 0.10±0.04 b |
| | V.d.+MTR17b | 80.95±9.91 ab | 57.05±6.52 ab | 19.05±6.73 ab | 14.85±2.03 bc | 0.40±0.18 ab |
| | V.d.+MTR17c | 76.19±9.52 ab | 49.66±7.35 ab | 23.81±6.15 ab | 11.73±2.35 c | 0.13±0.06 ab |
| | V.d.+TRIANUM-P | 52.38±6.74 b | 33.45±7.44 b | 4.76±4.76 b | 7.88±1.94 cd | 0.20±0.09 ab |

<sup>a</sup> Disease parameters were evaluated periodically on the basis of external symptoms during a period of 24 days (in Experiment I) and 30 days (in Experiment II) after root drenching with *Verticillium dahliae* conidial suspension (20 ml of $5 \times 10^6$ conidia ml<sup>-1</sup> per plant). One week prior to inoculation with *V. dahliae*, plants were root-drenched with bacterial suspension (20 ml of $10^8$ cfu ml<sup>-1</sup> of each strain per plant); whereas TRIANUM-P was also included in experiment II and applied by root drenching (20 ml of $3 \times$ $10^7$ cfu ml<sup>-1</sup> per plant). DI = final disease incidence; FDS = final disease severity; M = mortality; RAUDPC = relative area under the disease progress curve with reference to the maximum value potentially reached over each assessment period; IR = isolation ratio. Each value represents the mean of 21 plants after combining the results of 3 replicated experiments (except from IR that represents the mean of 5 plants in total). Within experiments, values in columns followed by the same letter are not significantly different according to Tukey's HSD test at $P \leq 0.05$

### Whole-Genome sequencing and analysis of selected endophytic bacterial isolates

Recently, whole-genome sequencing (WGS) of microbes has become more accessible and affordable as a tool for genotyping. Analysis of the entire microbial genome via WGS could provide unprecedented resolution in discriminating even highly related lineages of bacteria and revolutionize analysis that could help to understand their futures including antifungal and antibacterial activities [57, 58]. For this, we performed WGS of eleven selected isolates exhibiting interesting features, as well as, low 16S rDNA sequence similarity to known species. The genomes of these bacterial isolates were annotated using the RAST bacterial genome annotation tools (**Supplementary Figure S2**). Furthermore, all the genes related to the virulence, disease and defence that have been predicted at the particular isolates’ genomes, were identified (**Table 4** and **Table S2**). These data form the basis for future studies regarding the genome features that contribute to antifungal and antibacterial properties of these bacterial isolates.

**Table 4.** Number of genes related to Virulence, Disease and Defense features of the three new bacterial species identified in this study. The nalysis and the annotation was performed using the RAST genome annotation software.

| Virulence, Disease and Defense | *Arthrobacter* sp. CMR16 | *Pseudomonas* sp. CrR25 | *Pseudomonas* sp. CMR27 | *Pseudomonas* sp. CMR25 |
| --- | --- | --- | --- | --- |
| **Resistance to antibiotics and toxic compounds** | **19** | **56** | **42** | **45** |
| Mercury resistance operon | 1 | 0 | 0 | 0 |
| Copper homeostasis | 6 | 25 | 18 | 18 |
| Cobalt-zinc-cadmium resistance | 4 | 12 | 17 | 16 |
| Resistance to fluoroquinolones | 2 | 5 | 2 | 5 |
| Copper homeostasis: copper tolerance | 2 | 2 | 2 | 2 |
| Beta-lactamase | 1 | 0 | 2 | 1 |
| Mercuric reductase | 3 | 3 | 0 | 0 |
| Multidrug Resistance Efflux Pumps | 0 | 7 | 0 | 0 |
| Resistance to chromium compounds | 0 | 1 | 1 | 3 |
| **Invasion and intracellular resistance** | **19** | **21** | **14** | **17** |
| Mycobacterium virulence operon involved in protein synthesis (SSU ribosomal proteins) | 6 | 9 | 6 | 7 |
| Mycobacterium virulence operon involved in DNA transcription | 3 | 6 | 2 | 4 |
| Mycobacterium virulence operon possibly involved in quinolinate biosynthesis | 3 | 3 | 3 | 3 |
| Listeria surface proteins: Internalin-like proteins | 4 | 0 | 0 | 0 |
| Mycobacterium virulence operon involved in protein synthesis (LSU ribosomal proteins) | 3 | 3 | 3 | 3 |
| **Bacteriocins, ribosomally synthesized antibacterial peptides** | **0** | **2** | **2** | **2** |
| Tolerance to colicin E2 | 0 | 2 | 2 | 2 |
| **Membrane Transport** | **14** | **77** | **86** | **83** |
| Protein secretion system, Type II (Widespread colonization island) | 11 | 14 | 10 | 10 |
| Protein secretion system, Type II (General Secretion Pathway) | 0 | 15 | 0 | 0 |
| Protein secretion system, Type V (Two partner secretion pathway - TPS) | 0 | 4 | 0 | 0 |
| Protein secretion system, Type I | 0 | 0 | 29 | 22 |
| Protein secretion system, Type III | 0 | 0 | 0 | 0 |
| Protein secretion system, Type VI | 0 | 0 | 0 | 0 |
| Protein and nucleoprotein secretion system, Type IV (Type IV pilus) | 0 | 28 | 22 | 20 |
| Protein and nucleoprotein secretion system, Type IV (Conjugative transfer) | 0 | 12 | 0 | 0 |
| Protein secretion system, Type VII (Chaperone/Usher pathway, CU) | 0 | 0 | 13 | 12 |
| Twin-arginine translocation system | 3 | 4 | 7 | 7 |
| Protein secretion system, Type VIII (Extracellular nucleation/precipitation pathway, ENP) | 0 | 0 | 5 | 12 |

Interestingly, our whole-genome sequencing, the specific genes sequence comparison and the average nucleotide identity analysis, revealed the presence of three totally new bacterial species. To further investigate this finding we extracted the *recA* and *gyrB* gene sequences from the genomes and after concatenation we constructed phylogenetic trees using representatives from the related bacterial groups in order to investigate the phylogeny of our isolates (**Figures 6** **&** 7). Two of these species belong to *Pseudomonas* family, while the third is a member of *Arthrobacter* family. The isolates CMR25 and CMR27 seem to belong to an unidentified species of the *P. putida* group (**Figure 7**), and the isolate CrR25 appears to be an undefined species of the *P. mendocina* group (**Figure 7**). Their 16S rDNA sequence of both CMR25 and CMR27 are similar 99.8% to *Pseudomonas plecoglossicida* and the 16S rDNA of CrR25 is 98.75% similar to *Pseudomonas benzenivorans*. While the genome analysis and annotation the isolate CMR16 belong to an unidentified *Athrobacter* species (**Figure 6**), its 16S rDNA sequence assigns the isolate to *Paenarthrobacter nitroguajacolicus* with 98.78% similarity.

**Figure 6.**
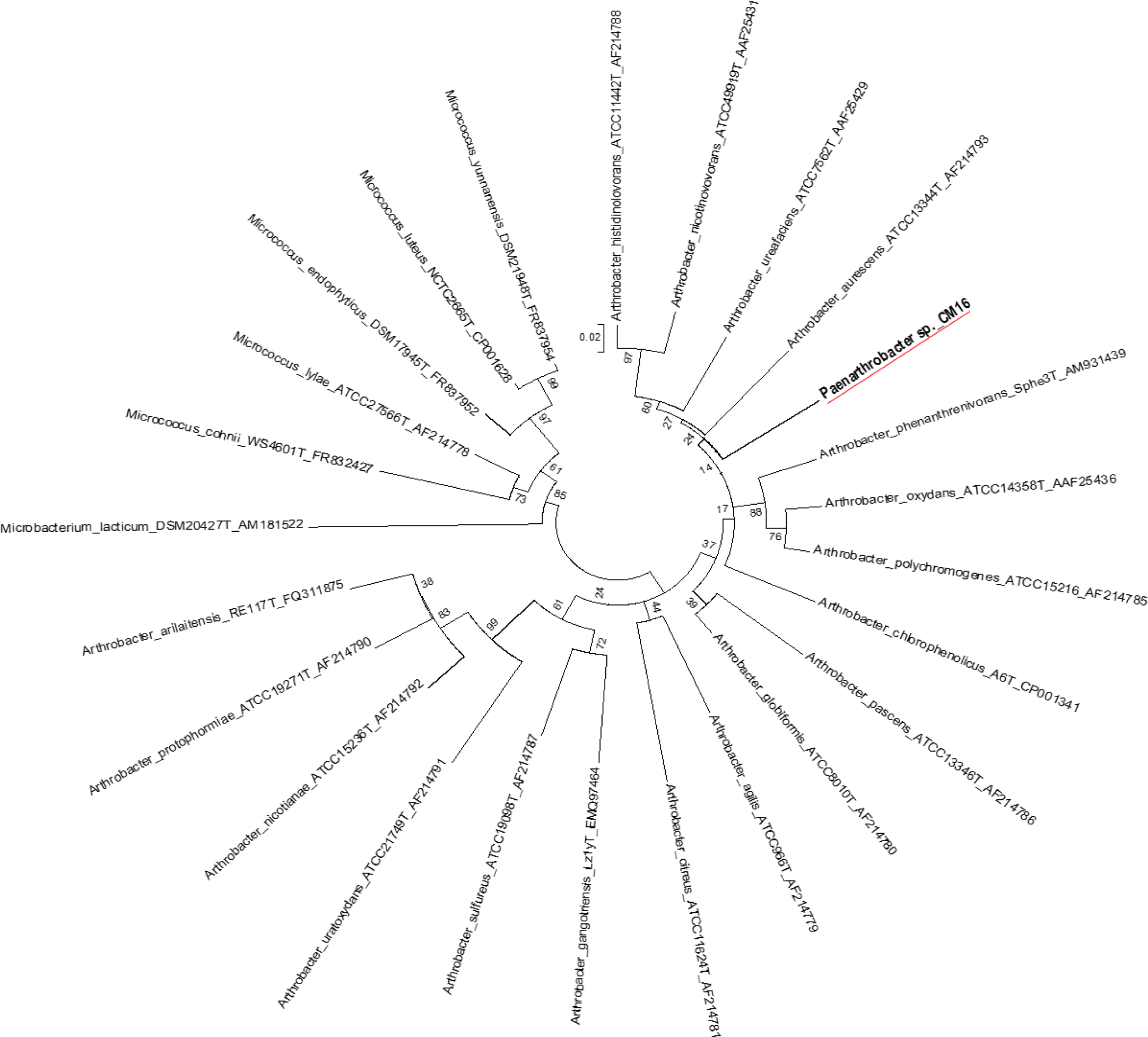
Molecular Phylogenetic analysis of *Arthrobacter recA-gyrB* genes by Maximum Likelihood method. The evolutionary history was inferred by using the Maximum Likelihood method based on the Tamura-Nei model [54]. The bootstrap consensus tree inferred from 500 replicates [55] is taken to represent the evolutionary history of the taxa analyzed [55]. Branches corresponding to partitions reproduced in less than 50% bootstrap replicates are collapsed. Initial tree(s) for the heuristic search were obtained automatically as follows. When the number of common sites was < 100 or less than one fourth of the total number of sites, the maximum parsimony method was used; otherwise BIONJ method with MCL distance matrix was used. The tree is drawn to scale, with branch lengths measured in the number of substitutions per site. The analysis involved 25 nucleotide sequences. Codon positions included were 1st+2nd+3rd+Noncoding. All positions with less than 95% site coverage were eliminated. That is, fewer than 5% alignment gaps, missing data, and ambiguous bases were allowed at any position. Evolutionary analyses were conducted in MEGA5 [34].

**Figure 7.**
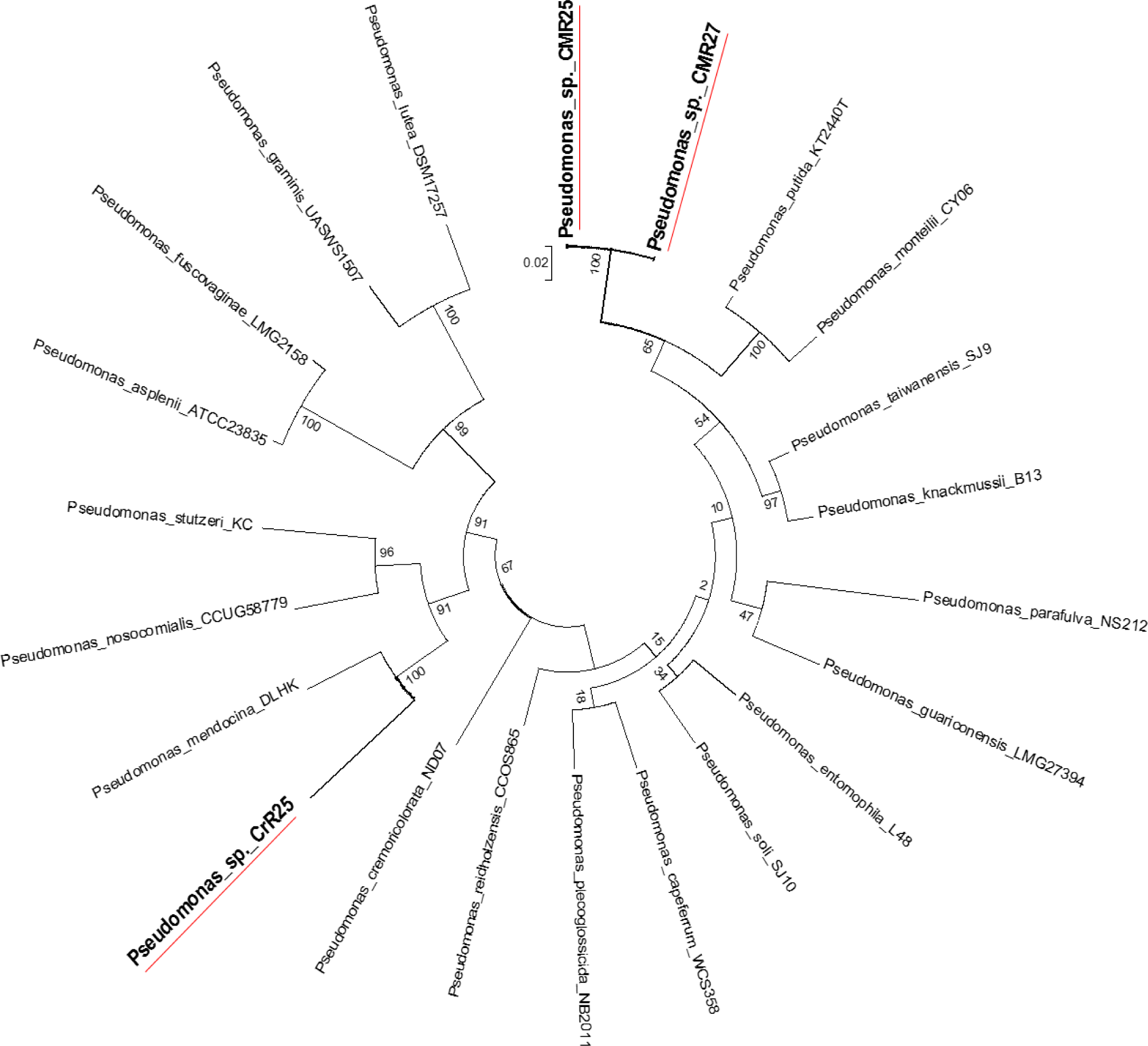
Molecular Phylogenetic analysis of the *recA-gyrB* genes from Pseudomonads belonging to *P. putida* and *P. mendocina* groups by Maximum Likelihood method. The evolutionary history was inferred by using the Maximum Likelihood method based on the Tamura-Nei model [54]. The tree with the highest log likelihood (-7831.6808) is shown. Initial tree(s) for the heuristic search were obtained automatically as follows. When the number of common sites was < 100 or less than one fourth of the total number of sites, the maximum parsimony method was used; otherwise BIONJ method with MCL distance matrix was used. The bootstrap consensus tree inferred from 500 replicates [55] is taken to represent the evolutionary history of the taxa analyzed [55]. The tree is drawn to scale, with branch lengths measured in the number of substitutions per site. The analysis involved 22 nucleotide sequences. Codon positions included were 1st+2nd+3rd+Noncoding. All positions with less than 95% site coverage were eliminated. That is, fewer than 5% alignment gaps, missing data, and ambiguous bases were allowed at any position. Evolutionary analyses were conducted in MEGA5 [34].

The identification of three new bacterial species in such a small number of WGS of our isolates, reveals the potentiality that the wild halophytic endophytome has regarding the isolation and the identification of new microbial species with novel capacities, that could be beneficial for both Agriculture (biotic and a-biotic stress tolerance, growth promotion, etc.) and potentially in clinical practice (identification of new antibiotics, antifungal compounds,

## Discussion

Of the nearly 391,000 species of vascular plants that exist in the earth, each is the host of a number of endophytes. Only a small number of these plants have ever been completely studied in respect to their endophytic composition. On the other hand, the use of endophytic microorganism for the control of both biotic and abiotic stresses, is relatively new and unexplored area of research. Endophytes have been studied for over two decades [35–37], but our understanding about the role of endophytic microbes in plant defense against biotic and abiotic stresses is still limited [38, 39]. The isolation, identification and the study of endophytes from plants that undergo continued abiotic stress could be essential for the development of proper biocontrol strategy for a sustainable agriculture and food security.

Even if scientific approaches on the diversity of endophytes have recently gained interest in plant-microbe research field, information on endophytes’ content and their potential contribution on the mutualistic interactions with cultivated plant species remain unclear. For the future precision in agriculture, targeted application of culturable beneficial microbial “artificial communities” may be a sustainable path to counteract biotic and abiotic stress conditions and ensure yield stability. However, within the endophytic bacterial communities, members show a strong influence on each other, which could include antagonistic, competitive, and mutualistic interactions [2]. This is important when introducing new species or communities into an agricultural field or when trying to isolate the causative beneficial species in complex microbiomes. Among the beneficial traits that are of particular significance, with potential to be exploited in crop ecosystems is the promotion of plant growth, the control of plant diseases and the high salinity tolerance. However, and apart from their use in agriculture, additional applications of endophytes could rely on their ability to produce a broad range of bioactive metabolites that are relevant for other purposes, including human health (antibiotics, antitumor compounds, etc.).

In this study, we investigate the abundances of taxa of the culturable bacterial endophytes of three halophytic plants, endemic in Crete Island, Greece, using culture-dependent techniques. We also investigate the proof-of-concept of using the halophytes as a valuable source of beneficial microbes that will be used in future agriculture. We tested our initial hypothesis that these endophytes can be used as potential biofertilizers and biocontrol agents for a sustainable agriculture. Taxonomically, all bacterial isolates belong to the phyla *Actinobacteria*, *Firmicutes* and *Proteobacteria* (either the *Alphaproteobacteria* or *Gammaproteobacteria* classes). While 24 different genera were identified, the three of the most abundant genera were *Bacillus* and *Enterobacter* and *Pseudomonas,* all of which have been previously observed before in studies of the endophytic microbiome of halophytes [16, 18, 40, 41].

*In planta* testing of *Oceanobacillus picturae* (CML15), *Terribacillus saccharophilus* (MTR05) and *Bacillus haikouensis* (CrR22) demonstrated an increase in both dry and fresh leaf weight in *Arabidopsis thaliana* plant under salinity stress. These strains are promising biofertilizers, since other strains of the same species have been shown to have plant growth promotion properties. *Terribacillus saccharophilus*, firstly reported at 2007, is a known halophilic bacterium that can grow on NaCl concentrations ranging 0-16% [42, 43]. This species is a known endophytic bacterium [44], that has been shown a systemic response that trigger an increase on monoterpenes, sesquiterpenes, tocopherols and membrane sterols, compounds engaged in antioxidant capacity in leaf tissues of grape resulting in stress tolerance [45]. *Bacillus haikouensis* is a recently described halotolerant bacterium isolated from paddy soil, able to grow on up to 17% NaCl concentrations [46]. *Oceanobacillus picturae* is a halophilic phosphate-solubilizing species with demonstrated siderophore production potential that has been isolated from saline environments and has been shown to promote plant growth promotion in mangroves and confer salinity stress tolerance in barley [47–49]. Many of our isolates showed ability to grow at high concentrations of salt (5% to 17% salinity), while it is important to mention that the Mediterranean Sea salinity is around 3.8%

Multiple strains that belong to the species *Bacillus licheniformis, Bacillus sonorensis, Pseudomonas glareae, Enterobacter hormaechei, Pseudomonas benzenivorans, Pseudomonas monteilii, Pseudomonas plecoglossicida* were shown to have strong antagonistic activity against the phytopathogenic bacteria *Ralstonia solanacearum* and *Clavibacter michiganensis* subsp*. michiganensis*, two very important plant pathogens with high economic impact on agriculture [50, 51]. Both are very important phytopathogens, since *Ralstonia* has a large host range able to infect more than 200 plant species easily adaptable in varying environmental conditions whereas *C. michiganensis* subsp. *michiganensis* is able to infect wheat, maize, potatoes and red and green peppers, despite its main host being tomatoes [51–53]. Moreover, a selected number of isolates that revealed *in-vitro* growth inhibition effect against *V. dahliae*, were also tested for their ability to inhibit *V. dahliae in-planta*. Several of these isolates revealed a very promising *in-planta* suppression effect of the polyphagous pathogen *V. dahliae*. Interestingly, isolates with strong *in-vitro* effect did not manage to inhibit *V. dahliae in-planta* but also isolates with medium or low *in-vitro* effect revealed very strong *in-planta *V. dahliae** growth inhibition. This data provide the proof of concept for our study but also indicate that in future studies we need to investigate all the isolates regarding their *in-planta* antifungal and/or antibacterial growth inhibition capacity.

Furthermore, the whole-genome sequencing (WGS) of a small number of our isolates revealed three new, unidentified bacterial species. The identification of three new species in a very small number isolates indicates the high potentiality that the wild halophytic endophytome has regarding the isolation and the identification of new microbial species with novel capacities, that could be beneficial for both Agriculture (biotic and a-biotic stress tolerance, growth promotion, etc.) and potentially in clinical practice (identification of new antibiotics, antifungal compounds, etc.).

The study of the endophytic microbes is now benefitting from a number of new culture methods and media, as well as, the rapid and efficient DNA based bacterial identification techniques and the total genome sequencing and annotation. The results of our study and the microbial collection we have generated, could be the basis for the future development of various synthetic “Bio-Inoculants”. They can also be the basis for a number of future studies, including the investigation of the colonization strategies that these microbes use, as well as, the elucidation of the molecular dialogs that take place during host-root colonization; the growth promotion; the salt tolerance and the immunity activation, by unique beneficial endophytes or artificial endophytic communities.

The endophytes must possess following attributes for agricultural exploitation: a) they must not be pathogenic and must not induce plant disease; b) they should be capable to spread and colonize the plant tissues; c) they should be culturable in order to be used in modern agriculture and d) they must colonize plant parts and to be compatible with the species genetic characteristics.

For a new environmentally friendly global strategy in food production that will be based in the sustainable agriculture with low chemical inputs and a low environmental impact, there is a strong need to search for novel beneficial entophytic microbes with as many desirable characters for enhancing the crop production. For this, the halophytic endophytes could be a environments, great source of beneficial microbes that could be a solution for agriculture in saline environments, as well as, a great source of microbes for biocontrol of important pathogens, based on biological systems involving CWR plants well-adapted to grow in soils affected by salinity and associated microorganisms are gaining interest.

## Acknowledgements

This project utilized equipment funded by the Wellcome Trust Institutional Strategic Support Fund (WT097835MF), Wellcome Trust Multi-User Equipment Award (WT101650MA) and BBSRC LOLA award (BB/K003240/1).

The authors acknowledge Dr. Karen Moore, Audrey Farbos, Georgina Morris of the Exeter Sequencing Service at University of Exeter, for DNA library preparation and genome sequencing.

This work benefitted from the support of the University of Exeter’s High-Performance

Computing (HPC) facility.

The project was partially supported by the Emblematic Action of the Greek General Secretariat for Research and Technology (GSRT), “AGRO4CRETE”, Protocol Number: SAE 013, Operational Program: SAE 013.

## Notes

### Competing Interest Statement

The authors have declared no competing interest.

