## Supplementary_Material for "Halophytic bacterial endophytome: a potential source of beneficial microbes for a sustainable agriculture"

**Supplementary Table S1.** Bacterial isolates from leaves and roots of *Matthiola tricuspidata*, *Crithmum maritimuma* and *Cakile maritima* plants and i*n vitro* salinity assays for salinity stress, phytopathogen growth inhibition and human pathogen growth inhibition.

**Supplementary_Table_S1_In_vitro_assays.xlsx**

**Supplementary Table S2.** Analysis of variance for disease incidence (DI), final disease severity (FDS), mortality (M), relative area under disease progress curve (RAUDPC) and pathogen isolation ration (IR) for eggplants artificially inoculated with V. dahliae and treated with various bacterial strains and TRIANUM-P (CrR14, CrR18, CrR4, MTR12, MTR18, CM1, CM3, CM4, CM25 in *experiment I*, and MTR17a, MTR17d, MTR17f, MTR17g, MTR17h and MTR17b, MTR17c, TRIANUM-P in *experiment II*) or not (C-, V.d.).

| | | ***F* values**[a] | | | | |
|---|---|---|---|---|---|---|
| | | **Experiment I** | | | | |
| **Source** | **df**[b] | **DI** | **FDS** | **M** | **RAUDPC** | **IR** |
| Replication | 2 | 0.57 | 0.02 | 0.50 | 0.67 | 1.82 |
| Treatment | 10 | 362.38*** | 96.82*** | 7.65*** | 31.57*** | 2.84* |
| Replication × Treatment | 20 | 0.57 | 1.10 | 0.42 | 1.23 | 0.92 |
| | | **Experiment II** | | | | |
| **Source** | **df**[b] | **DI** | **FDS** | **M** | **RAUDPC** | **IR** |
| Replication | 2 | 0.72 | 0.52 | 0.74 | 1.77 | 0.12 |
| Treatment | 10 | 9.58*** | 12.05*** | 3.27** | 11.57*** | 2.83* |
| Replication × Treatment | 20 | 1.13 | 1.09 | 1.20 | 1.19 | 0.71 |

[a] Symbols '*' and '***' indicate significance at $P \leq 0.05$ and 0.001 levels, respectively, according to the *F* test

[b] degrees of freedom between groups

**Supplementary Table S3.** Analysis of variance for plant height, plant fresh weight and total number of leaves for eggplant artificially inoculated with *V. dahliae* and treated with various bacterial strains and TRIANUM-P (CrR14, CrR18, CrR4, MTR12, MTR18, CM1, CM3, CM4, CM25 in experiment I, and MTR17a, MTR17d, MTR17f, MTR17g, MTR17h, MTR17b, MTR17c, TRIANUM-P in *experiment II*) or not (C-, V.d.).

| | | ***F* values**[a] | | |
|---|---|---|---|---|
| | | **Experiment I** | | |
| **Source** | **df**[b] | **Height** | **Weight** | **Leaves** |
| Replication | 2 | 2.43 | 1.12 | 1.65 |
| Treatment | 10 | 17.09*** | 111.03*** | 12.23*** |
| Replication × Treatment | 20 | 1.44 | 0.47 | 1.34 |
| | | **Experiment II** | | |
| **Source** | **df**[b] | **Height** | **Weight** | **Leaves** |
| Replication | 2 | 1.58 | 1.05 | 0.56 |
| Treatment | 10 | 3.26*** | 12.60*** | 1.42 |
| Replication × Treatment | 20 | 1.22 | 1.21 | 0.87 |

[a] Symbol '***' indicates significance at $P \leq$ 0.05 and 0.001 levels, respectively, according to the *F* test

[b] degrees of freedom between groups.

**Supplementary Table S4.** Bacterial genome features in the categories of virulence, disease and defence for bacterial isolates CMR4, CrR16, CrR7, CMR29, CrR6, CrR14 and CrR18.

**Supplementary_Table_S4_Bacterial Genomes features_.xlsx**

**Supplementary Figure S1.** Plant collection sites on Crete Island**.**

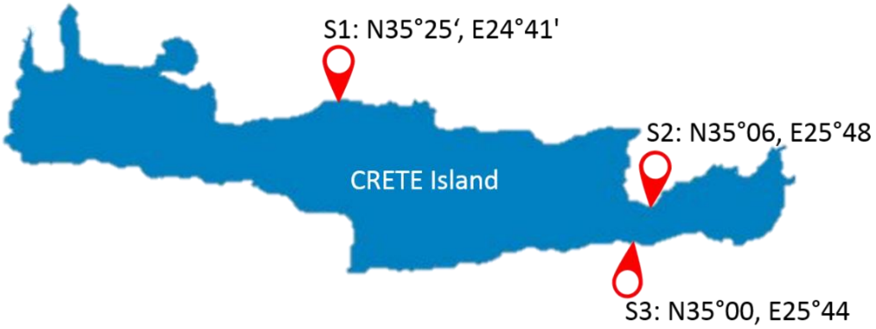

**Supplementary Figure S2.** Subsystem feature counts, category distribution and subsystem coverage of the gene content of fully sequenced bacterial isolates.

### *Pseudomonas seleniipraecipitans* CrR14

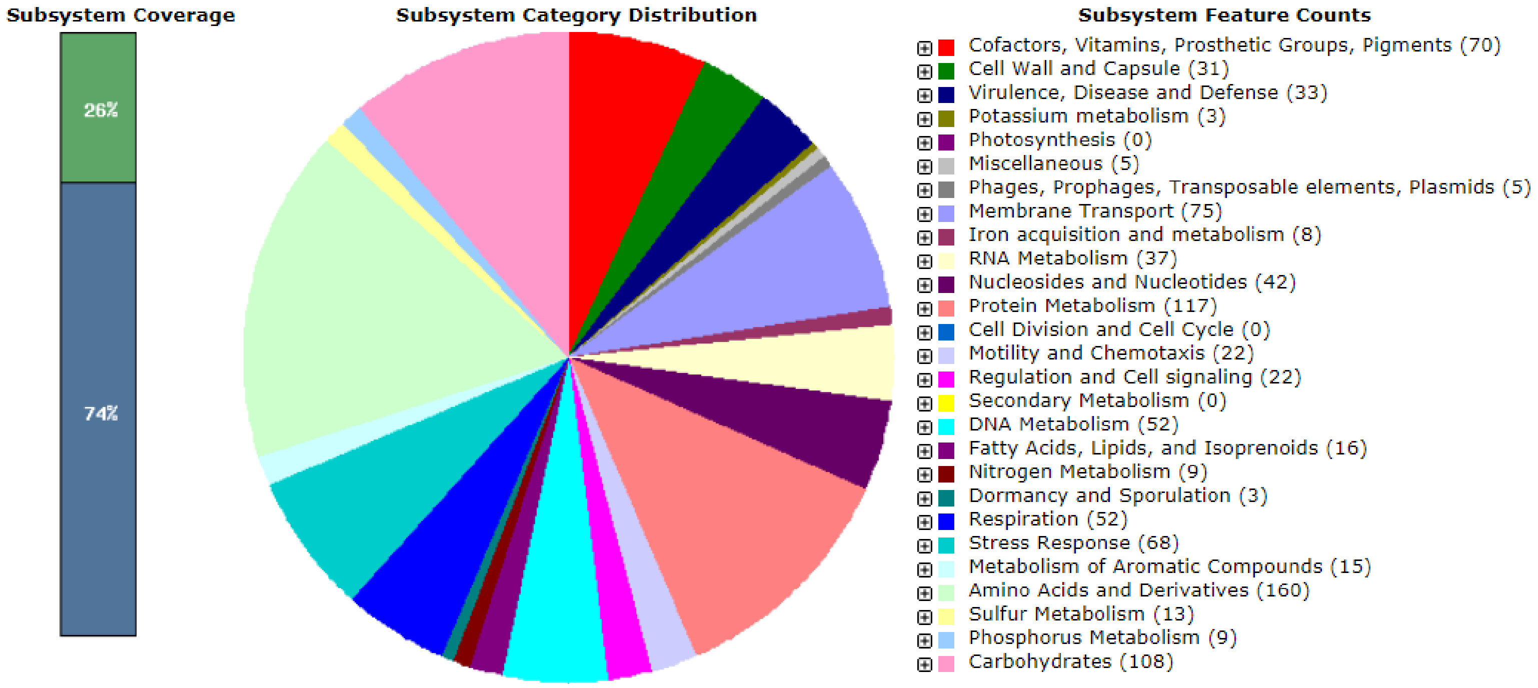


### *Bacillus altitudinis* CML04

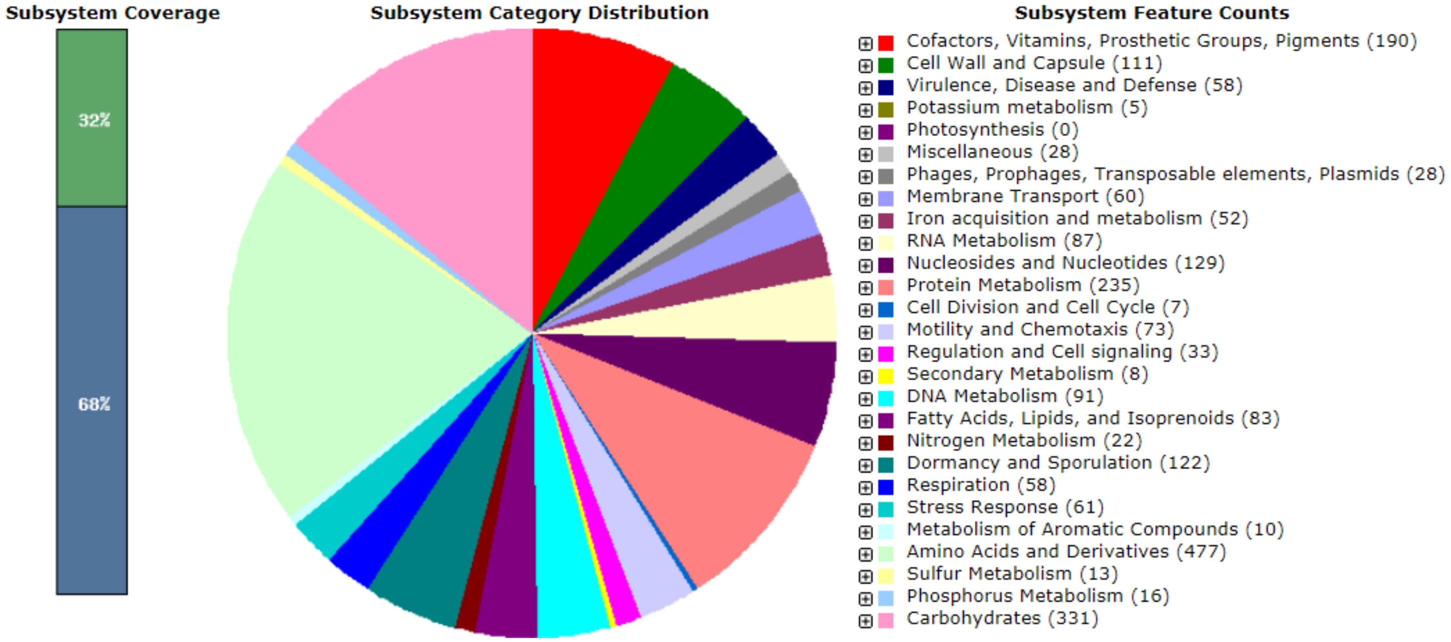

### *Bacillus haikouensis* CrR16

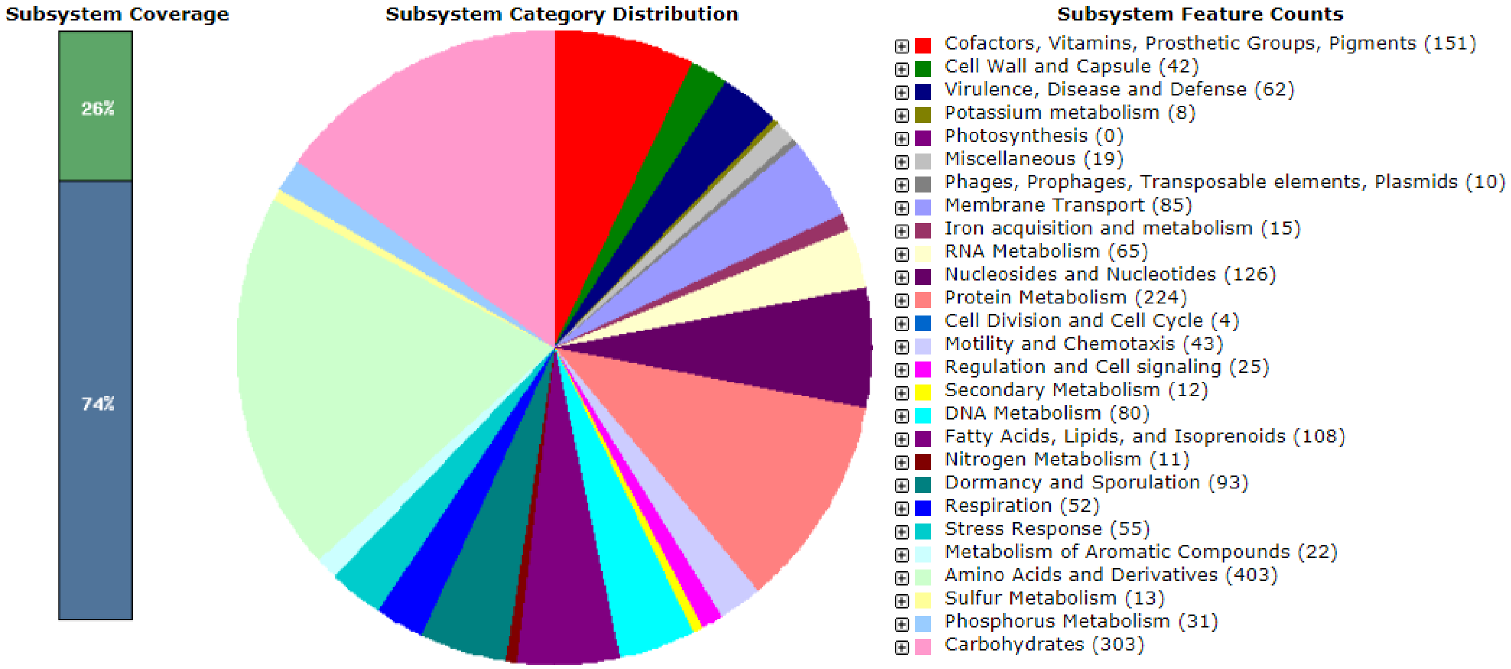


### *Enterobacter hormaechei* CrR07

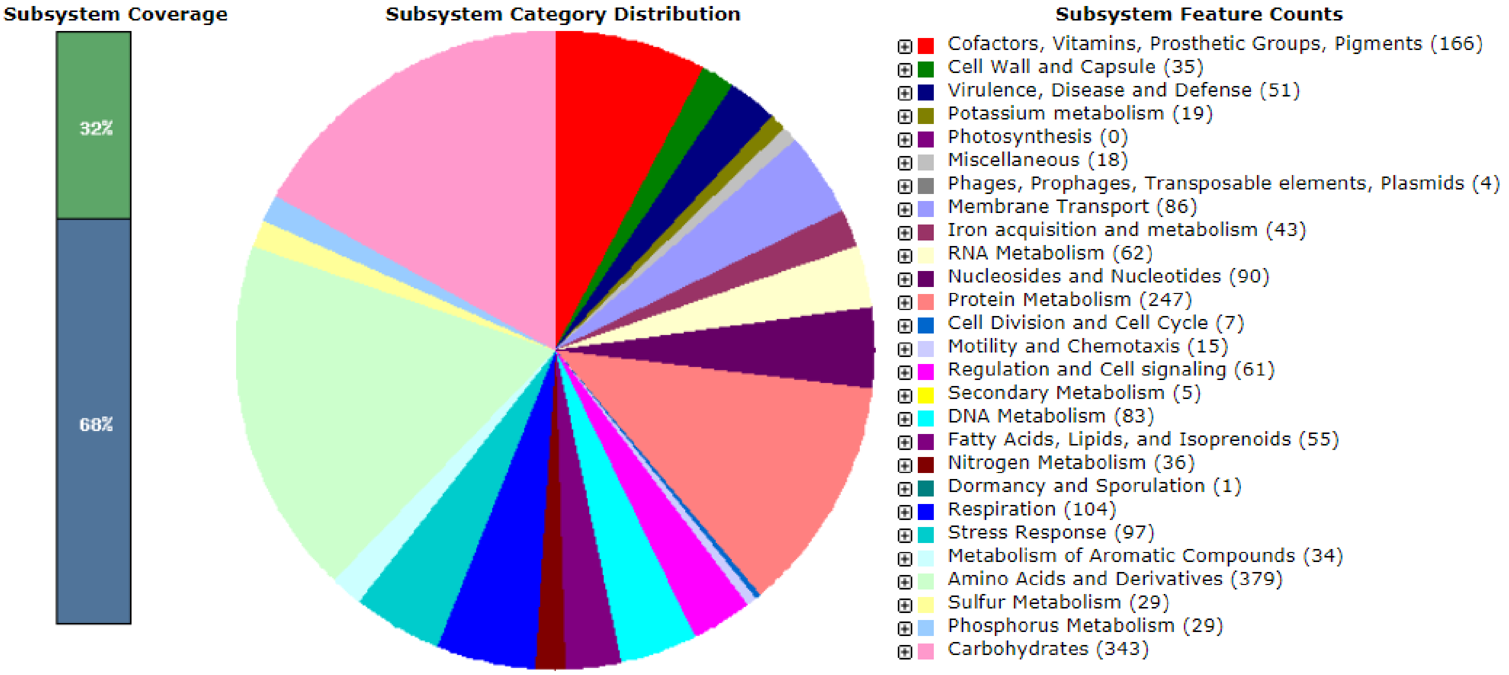

***Enterobacter hormaechei* CMR29**

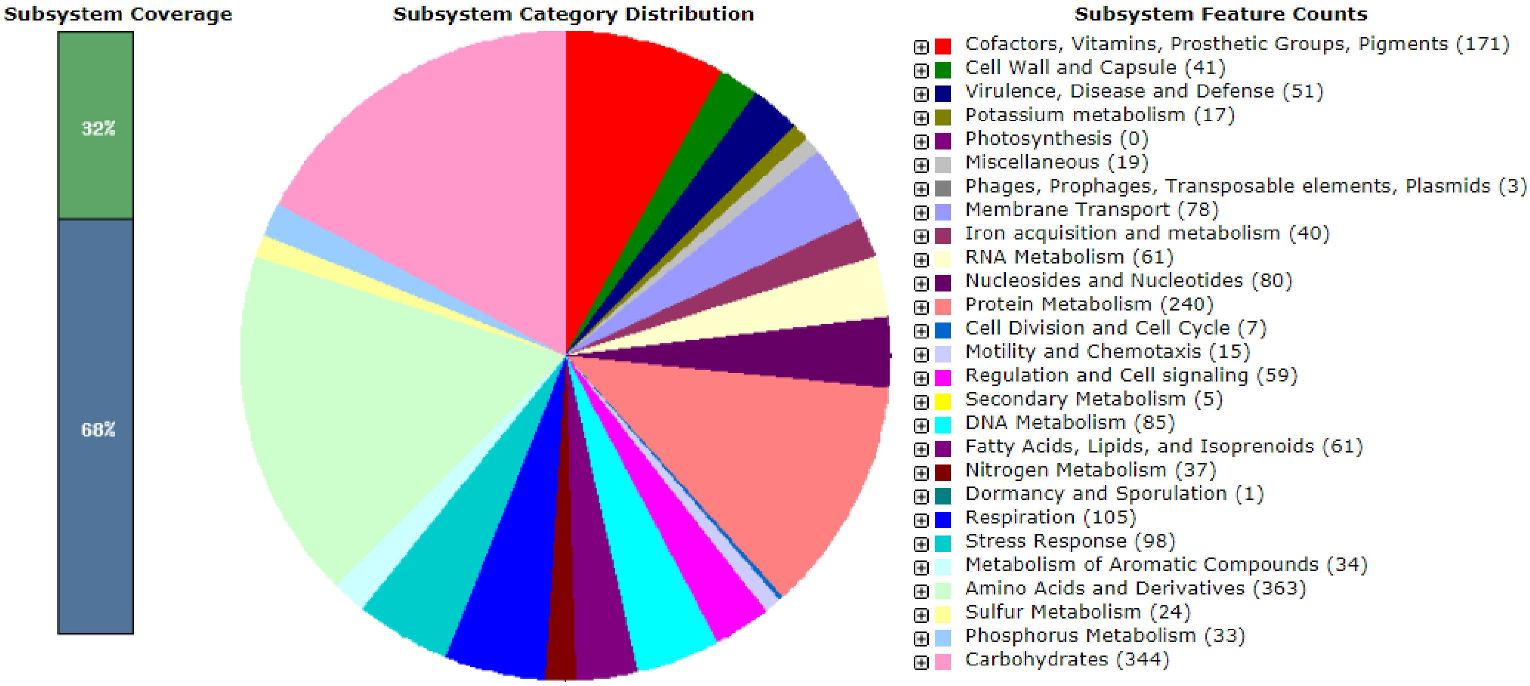


***Glutamicibacter halophytocola* CrR06**

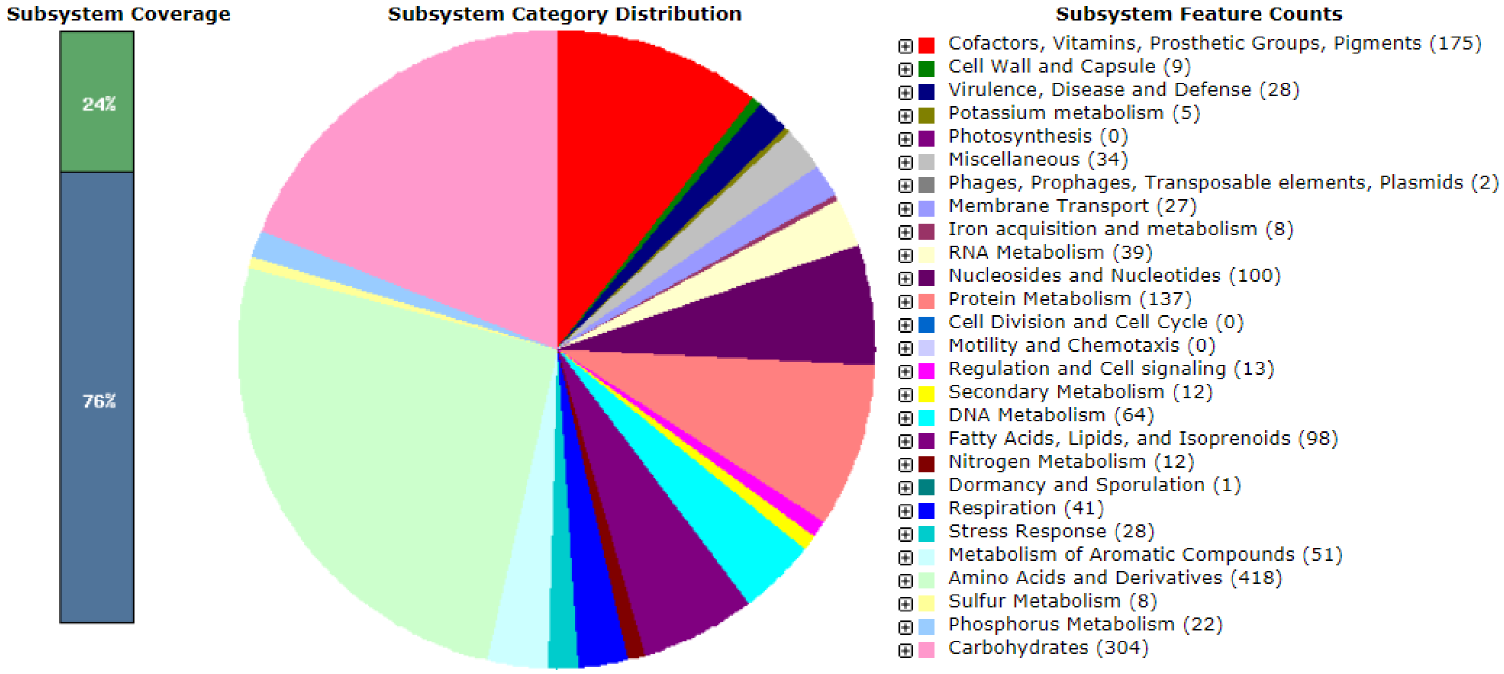

### *Sphingobacterium shayense* CrR18

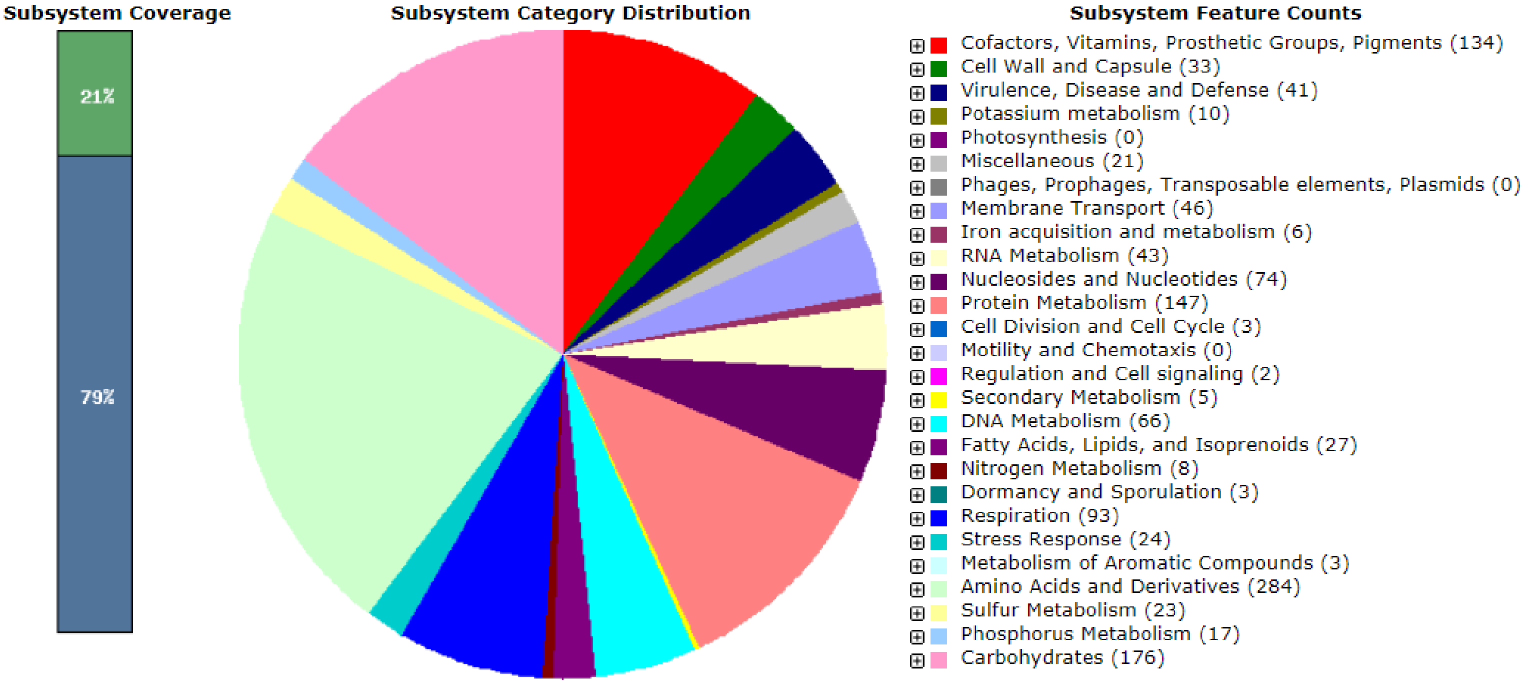


### *Arthrobacter* sp. CMR16

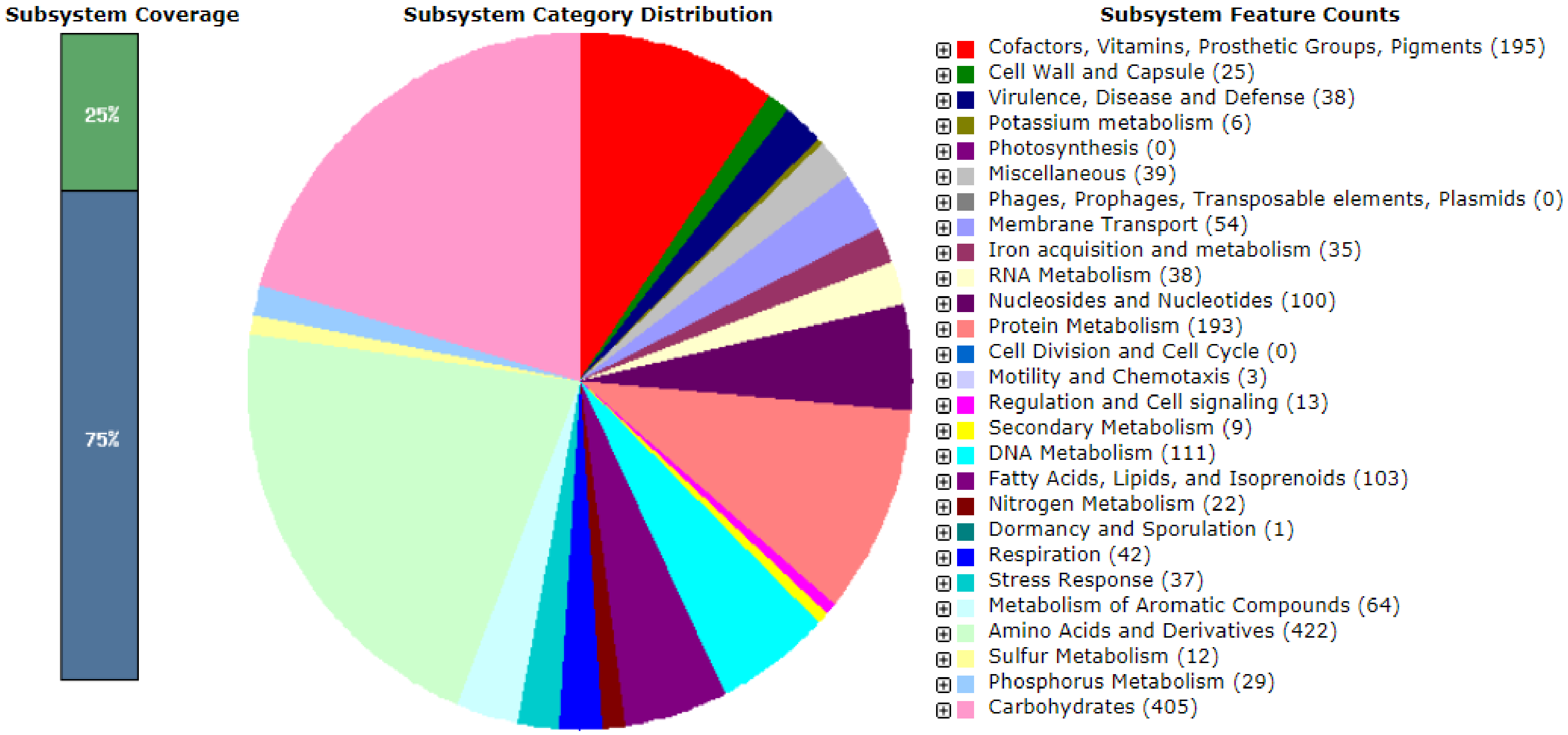

### *Pseudomonas* sp. CrR25

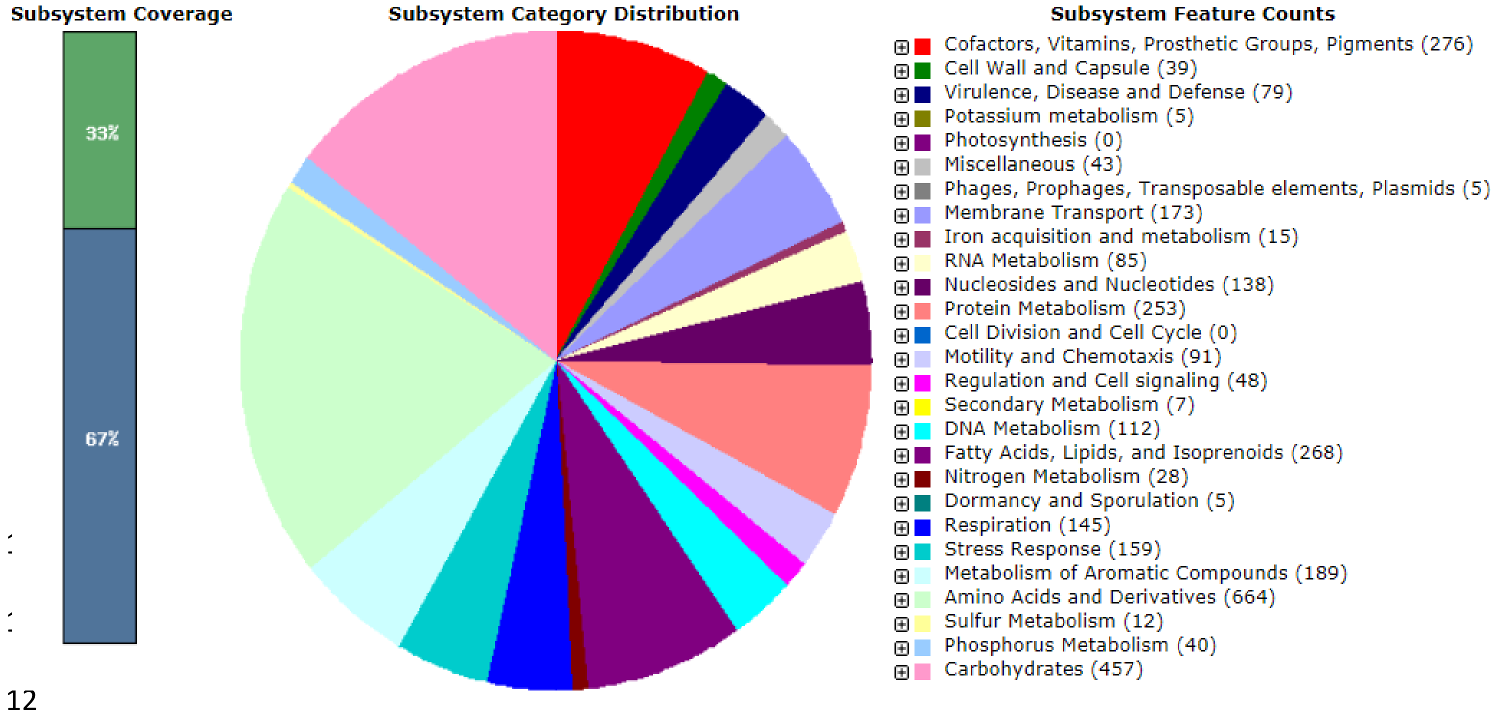


### *Pseudomonas* sp. CMR27

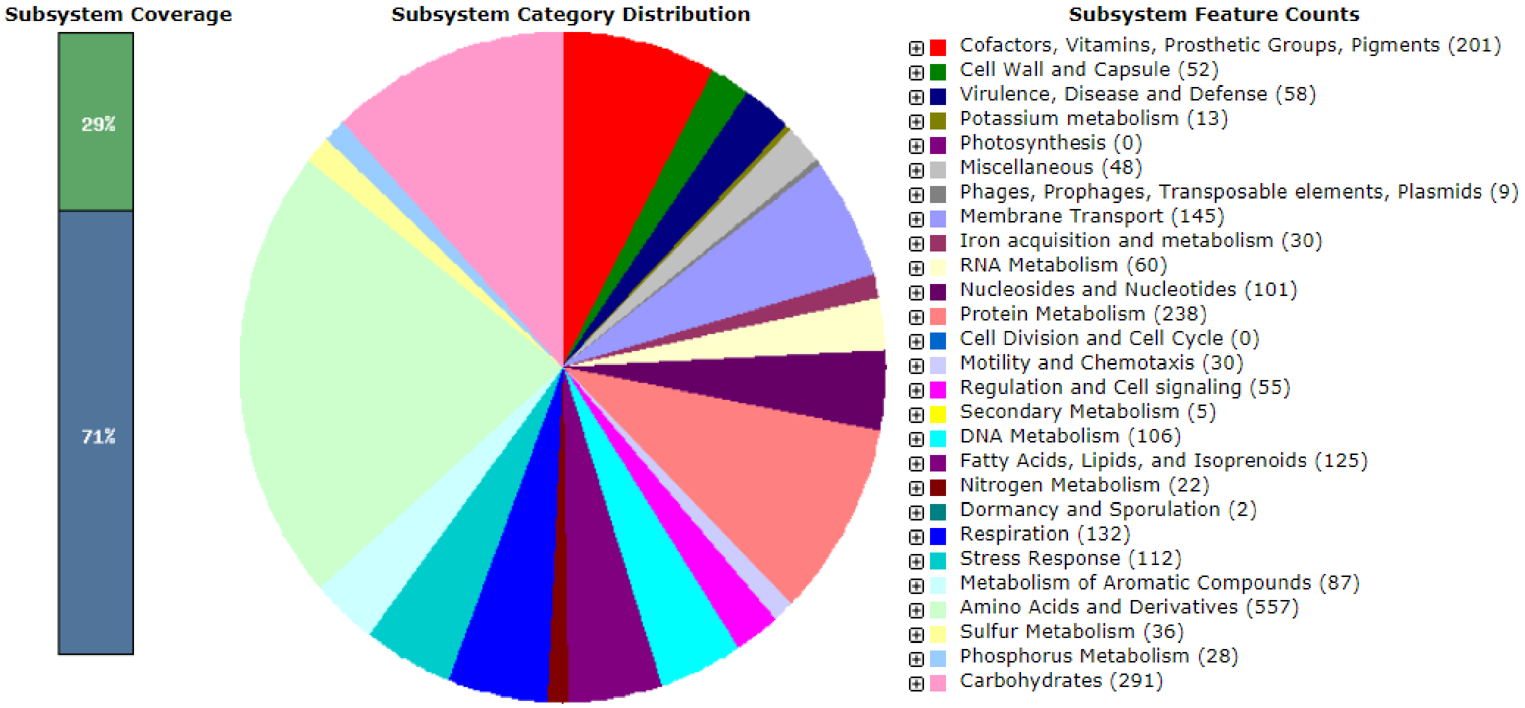

*Pseudomonas* sp. CMR25

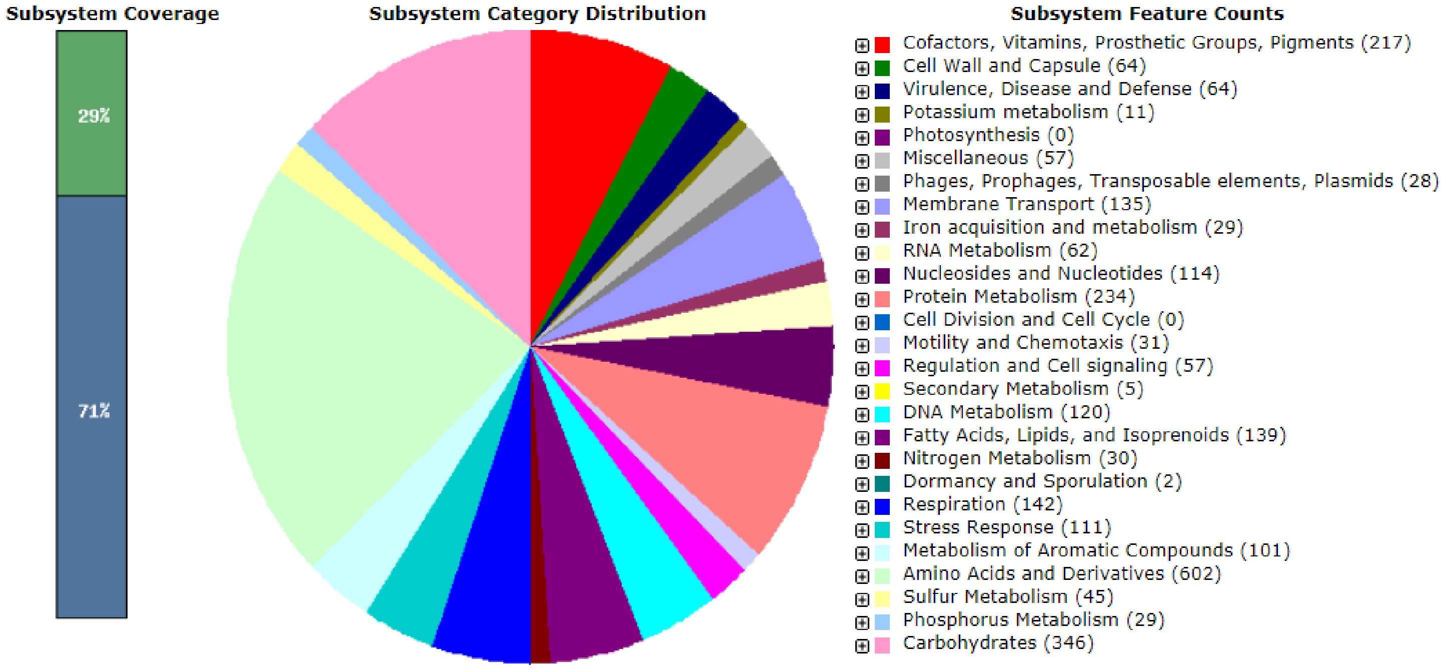
